# Systems-level analysis identifies protein-RNA interactions contributing to autotrophy of *Clostridium autoethanogenum*

**DOI:** 10.64898/2025.12.18.695208

**Authors:** Angela Re, Gianfranco Michele Maria Politano, Kristina Reinmets, Alfredo Benso, Laurence Girbal, Muriel Cocaign-Bousquet, Kaspar Valgepea

## Abstract

Burning fossil fuels drives climate change with detrimental consequences on humankind while waste accumulation from increasing consumption threatens biosustainability globally. Gas fermentation enables to recycle carbon oxides (CO and CO_2_) from industrial waste gases and gasified waste into value-added products using gas-fermenting microbes, namely acetogens. However, our limited understanding of gene function and metabolic regulation is hindering rational engineering of acetogen cell factories. In this work, we identified, for the first time in acetogens, genome-wide protein-RNA interactions contributing to autotrophy in the model-acetogen *Clostridium autoethanogenum* by combining steady-state chemostat cultivation, functional genomics, and computational methods. We first detected limited and uncoupled transcriptional and translational regulation between autotrophy and heterotrophy. Rigorous mapping of genome-wide transcriptional architecture revealed both differential usage and signal strength of transcriptional start and termination sites between genes and growth substrates. We then used computational tools to reconstruct protein-RNA interactions for differentially regulated genes predicting 14 trans-acting regulatory RNA-binding proteins (RBPs) involved in post-transcriptional regulation and contributing to autotrophy. Most RBPs, two of which are translationally regulated, perform RNA modifications and regulate mRNA stability while others target translation-related genes. Our work provides valuable knowledge for metabolic engineering of acetogens and potentially contributes towards understanding primordial life on Earth.

## Introduction

Our continued reliance on fossil fuels drives greenhouse gas emissions that are accelerating climate change with detrimental consequences on the environment and humankind. In addition, increasing accumulation of waste from increasing global population and prosperity is threatening biosustainability through degradation of ecosystems. Gas fermentation has emerged as an attractive technology to tackle these challenges by recycling waste carbon from industrial off-gases and gasified waste into value-added products using gas-fermenting microbes^1,2^. Acetogens are the preferred biocatalysts for capture and conversion of carbon oxides (CO and CO_2_ ) into fuels and chemicals^3,4^ as they can use gas as their sole carbon and energy source^5^ by employing the most-efficient CO_2_ -fixation pathway known to date^6,7^, the Wood-Ljungdahl pathway (WLP)^5,8^. However, our limited understanding of gene function and metabolic regulation in acetogens is hindering rational engineering of acetogen cell factories^3,4,9-11^.

Most studies aimed at elucidating the relationship between genotypes and phenotypes in acetogens have focused on transcription^3^. Results from a limited set of experimental conditions suggest that metabolic fluxes in acetogens are primarily regulated post-translationally^12-14^. However, comprehensive analysis of acetogen multi-omics datasets^15-18^ highlights that post-transcriptional regulation of gene expression has a significant effect on the flow of genetic information from DNA to protein^19^, in a similar way to other microbes^20^. Key players in post-transcriptional regulation in microbes are RNA-binding proteins (RBPs)^20^, encompassing direct^21^ or indirect^22^ regulation of transcription^23-25^, RNA turnover^26^, and translation^23,27-29^. Additionally, RBPs can play key roles in regulating microbial metabolism through distinct RBP-RNA interactions^30,31^. However, no RBPs have been described in acetogens so far, and thus we lack knowledge about RBP-RNA interactions. Hence, fundamental understanding of acetogen metabolism and potentially also engineering of cell factories could be advanced through uncovering the unexplored space of RBPs.

The aim of our work was to identify, for the first time in acetogens, genome-wide RPB-RNA interactions that contribute to autotrophy. We used the model-acetogen *Clostridium autoethanogenum* as its metabolism has been explored through numerous omics datasets^3,32-37^, it has been engineered to produce various non-native chemicals^3,38,39^, and it is used in commercial-scale gas fermentation for ethanol production^2^. We firstly aimed to experimentally determine RBPs using the RNA interactome capture (RIC) technique^40,41^ that was recently adapted for the microbe *Escherichia coli*^4^, but after numerous attempts failed to engineer a *C. autoethanogenum* strain expressing a heterologous poly(A) polymerase that is needed for genome-wide RNA polyadenylation and pull-down in RIC. We thus employed an alternative comprehensive approach to predict RBPs and RBP-RNA interactions by combining functional genomics and computational methods (**Fig. 1**).

**Fig. 1.**
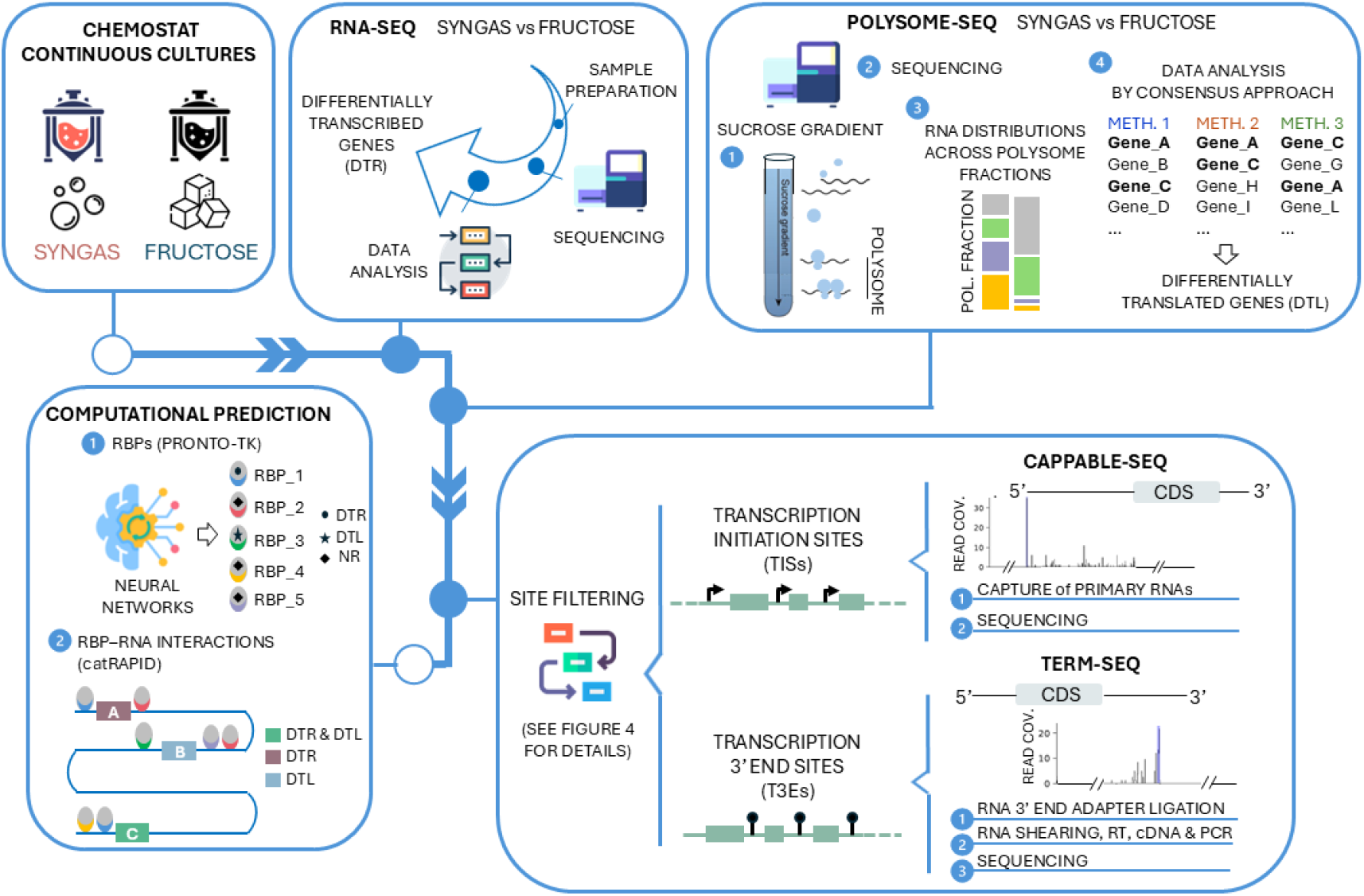
Integrative functional genomics and computational approach used in this work to predict RNA-binding proteins (RBPs) and protein-RNA interactions contributing to autotrophy of *C. autoethanogenum*. CDS, coding DNA sequence; DTR, differentially transcribed; DTL, differentially translated; NR, not differentially regulated.

Our approach first quantified both genome-wide transcriptional and translational regulation to identify transcripts differentially transcribed and/or translated during autotrophy vs heterotrophy. Transcription levels were quantified using RNA-seq while polysome-seq instead of Ribo-seq was used to evaluate translational status of transcripts since only the former offers resolution at the level of each mRNA copy^43^. Next, we quantified genome-wide transcriptional architecture by accurate mapping of 5’-untranslated regions (UTR) and 3’UTRs at single nucleotide resolution to determine the RNA sequences where RBPs most likely bind^44,45^. 5’UTRs were determined through identification of transcription initiation sites (TISs) by Cappable-seq^46^, which simplifies data generation and analysis, and improves reproducibility and robustness^46,47^ over the widely used alternative method dRNA-seq^48^. 3’UTRs were identified by mapping transcription 3’ ends (T3E) using Term-seq^49^ that has an advantage of ensuring that the sequences adjacent to adapters in the sequencing reads represent RNA 3’ termini originally present in the samples. We then used PRONTO-TK^50^, a neural network-based toolkit, to identify potential RBPs in *C. autoethanogenum* as it uses ProtT5-XL protein embeddings and deep learning to predict GO term-based functions with high precision and computational efficiency. Finally, high-resolution transcript boundaries allowed us to computationally reconstruct RBP-RNA interactions contributing to autotrophy in *C. autoethanogenum* using the inferential tool *cat*RAPID^51^, which uniquely combines high-throughput prediction of protein-RNA interactions with multi-parametric rating in an organism-agnostic framework. Importantly, all these analyses were performed from the same autotrophic and heterotrophic cultures that were grown in bioreactor chemostat cultures, thus yielding steady-state data that enable quantification of strictly defined and condition-dependent phenotypes^52,53^.

While our work detected limited and uncoupled transcriptional and translational regulation between autotrophy and heterotrophy, *C. autoethanogenum* showed both differential usage and signal strength of transcriptional start and termination sites between genes and growth substrates. Reconstruction of protein-RNA interactions identified 14 trans-acting regulatory RBPs involved in post-transcriptional regulation of differentially regulated genes for autotrophy. Our work provides valuable knowledge and potential targets for metabolic engineering of acetogens and potentially contributes towards understanding primordial life as the WLP could be first biochemical pathway on Earth^54^.

## Results

### Autotrophic and heterotrophic chemostat cultures of C. autoethanogenum

To map genome-wide RPB-RNA interactions involved in enabling autotrophy in acetogens, we grew the *C. autoethanogenum* strain LAbrini^35^ on syngas (CO, H_2_, CO_2_ ) and fructose in chemostat continuous cultures (**Fig. 1**). Importantly, this allowed us to grow cells at the same specific growth rate (µ; ∼0.04 h^-1^ or ∼1 day^-1^), as µ equals dilution rate at steady-state, and thus facilitated unequivocal quantification of transcriptional and translational patterns of autotrophy and heterotrophy without the confounding effects from different µ during exponential autotrophic and heterotrophic batch growth. For instance, from the 2,030 differentially expressed transcripts between *C. autoethanogenum* syngas and fructose batch cultures^55^, ∼24% were also differentially expressed with faster growth in autotrophic chemostat cultures^33^.

Notably, we found that both specific acetate and ethanol production rates (mmol per gram of dry cell weight per day) were ∼2.3-fold higher and the specific 2,3-butanediol production rate was 9-fold higher for syngas vs fructose steady-state cultures (**Supplementary Fig. 1a**). The former can be explained by a doubled allocation of carbon into biomass (**Supplementary Fig. 1b**) that is favoured by the more energy-rich nutrient fructose while the latter shows the higher need to regenerate reducing equivalents with by-products during autotrophy when less biomass can be produced. Interestingly, both of our chemostat cultures with the LAbrini strain showed only ∼1.5-fold higher acetate production than ethanol (**Supplementary Fig. 1a**) while nearly no ethanol production was detected in fructose-grown batch cultures of *C. autoethanogenum* strain JA1-1^55^.

### Limited transcriptional and translational control involved in realising autotrophy

We first quantified transcriptional and translational regulation involved in autotrophy (**Fig. 1**) using RNA-seq (**Supplementary Table 1**) and polysome-seq (**Supplementary Table 2**). We note that polysome profiling has been performed only for a few microbes^43,56-59^ but not for Clostridia or acetogens. While our RNA-seq analysis quantified transcript levels for 3,557 genes, polysome-seq analysis determined the translation status for 3,875 genes on average across bio-replicates of syngas and fructose cultures. The reproducibility of bio-replicates across syngas and fructose cultures was estimated with Spearman correlation coefficients of 0.99 and 0.71 for RNA-seq and polysome-seq data (**Supplementary Fig. 2a and 3**), respectively. Bio-replicates for RNA-seq data also clustered well showing clearly distinguished transcriptome profiles between syngas and fructose cultures (**Supplementary Fig. 2b and 2c**). Analysis of polysome-seq data (see Methods for details) led to the removal of an outlier bio-replicate culture for both syngas and fructose (**Supplementary Fig. 4**). Analysis of RNA-seq data yielded 1,784 genes with statistically differential expression (q-value < 0.05) between syngas and fructose growth (**Fig. 2a; Supplementary Table 3**). We determined the top 5% ranking ones according to q-value (lower value ranks higher) as differentially transcribed genes (DTRs) to ensure consistency with determination of differentially translated genes (DTLs) from polysome-seq data (see next paragraph). Thus, 176 genes were considered as DTR with 109 and 67 being down-or up-regulated, respectively, on syngas vs fructose (**Fig. 2a; Supplementary Table 3**).

**Fig. 2.**
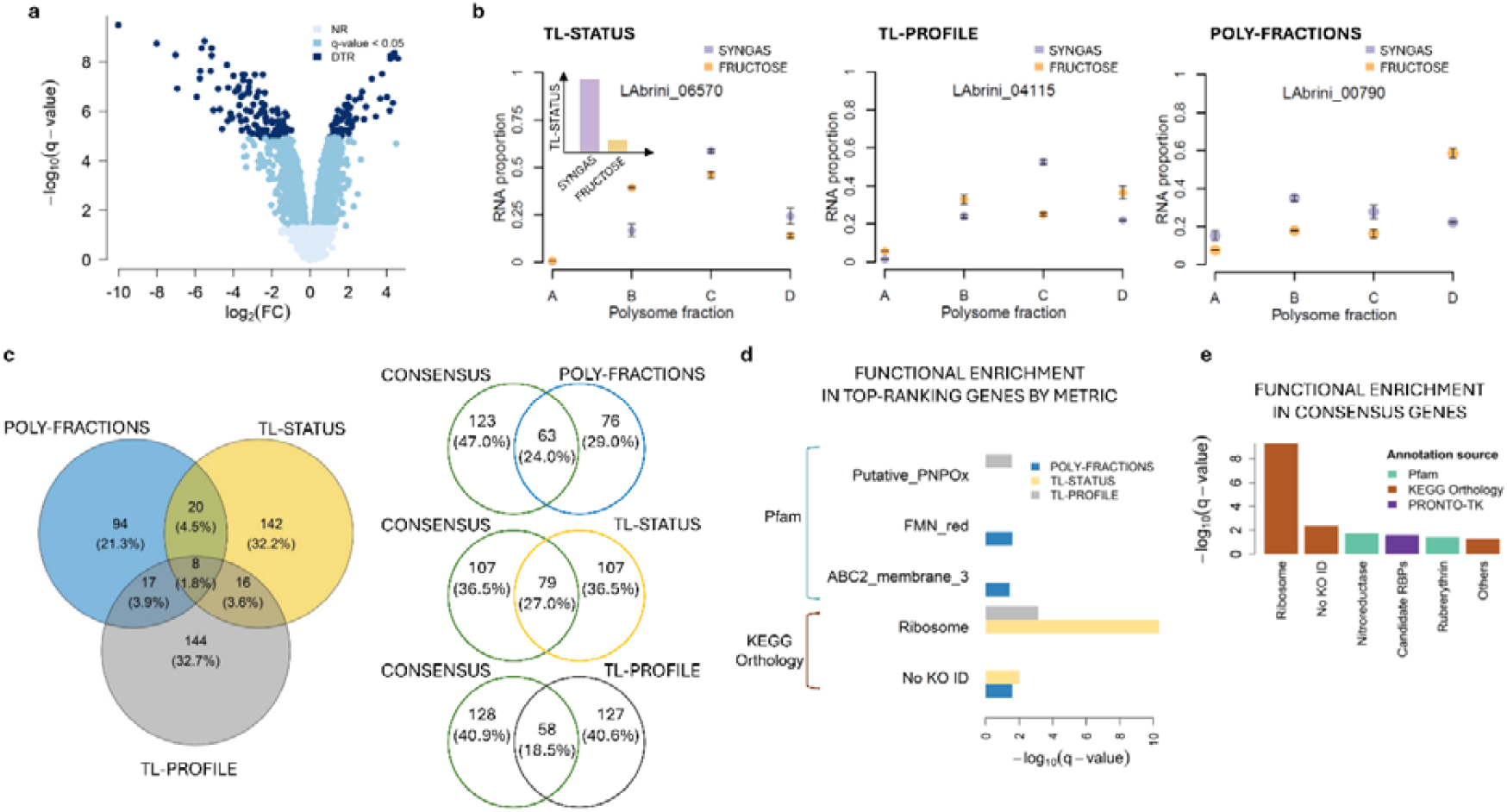
Limited transcriptional and translational control of autotrophy in *C. autoethanogenum*. **a** Volcano plot showing genes determined as differentially transcribed (DTR; #176) as a subset of 1,784 genes with statistically different expression (q-value < 0.05) between syngas and fructose chemostats. NR, not differentially regulated; FC, fold-change **b** Representative polysome profiles of top-ranking genes for each of the three metrics display the mRNA proportions of a gene across the ribosome-free fraction (A) and the polysome-bound fractions B (monosome), C (low ribosome density), and D (high ribosome density). For the TL-STATUS metric, inset shows the calculated translation status for syngas and fructose chemostats. Data are average ± standard deviation between biological duplicate chemostat cultures. **c** The three-set Venn diagram showing the overlaps between the differentially translated genes (DTL) identified by the POLY-FRACTIONS, TL-STATUS, and TL-PROFILE metrics. The two-set Venn diagrams show the overlap between the DTL identified by the CONSENSUS approach and each of the three metrics. **d** Functional classes enriched in DTLs according to each metric by hypergeometric test (FDR = 5%). Pfam, Pfam entries^61^ from eggNOG-mapper 2.1.12^62^ ; KEGG Orthology, KEGG Orthology (KO) functional categories^60^ from Valgepea *et al*. (Level 3 in Table S3^13^ ); NS, not significant. **e** Functional classes enriched in DTLs according to CONSENSUS approach by hypergeometric test (FDR = 5%). PRONTO-TK, computationally predicted RNA-binding proteins by PRONTO-TK^50^ .

Since polysome-seq quantifies transcripts bound to different numbers of ribosomes at single mRNA copy level, it allows to capture distinct traits compared to Ribo-seq that are indicative of translational control. To take advantage of this for determining DTLs, we established three complementary metrics for translational regulation (**Fig. 2b**; see Methods for details). First, we quantified the difference in translation status per gene between syngas and fructose (TL-STATUS), where the translation status is calculated as the ratio between the sum of the mRNA proportions quantified in the ribosome-bound and ribosome-free fractions. Positive value for TL-STATUS indicates up-regulation and negative down-regulation on syngas vs fructose (**Supplementary Table 4**). The translation status defined here is equivalent to the term ‘ribosome occupancy’ traditionally used for polysome-seq data^43^. Our data show that >90% of mRNA copies for each gene are ribosome-bound (**Supplementary Table 2**), similar to *E. coli*^58^ but higher than in *Lactococcus lactis*^59^ and *Saccharomyces cerevisiae*^43,57^. Second, we quantified the difference between the polysome profiles in the two growth conditions for each gene using the variance-weighted Aitchison’s distance (TL-PROFILE) (**Supplementary Table 4**). Third, we quantified for each gene the number of polysome fractions for which the difference in mRNA proportions between syngas and fructose was statistically significant by bootstrapping (POLY-FRACTIONS) (**Supplementary Table 4**).

To evaluate these three metrics as indicators of gene-level translational regulation involved in realising autotrophy, we compared the genes within the top 5% ranks of each metric. From the total of 441 genes among the top 5% ranks, ∼2% (8) and ∼14% (61) were shared between three or two metrics, respectively (**Fig. 2c**). Interestingly, each metric identified on average ∼29% unique genes demonstrating the relevance of using multiple metrics to evaluate translational regulation. Consistent with this was the enrichment of unique KEGG Orthology functional categories^60^ or Pfam entries^61^ within the top 5% gene sets of each metric (**Fig. 2d**). Therefore, to capture the potentially unique information on translational regulation carried by the different metrics while also considering the general overlap between the metrics, we decided to determine DTLs from polysome-seq data as the top 5% genes from ranking the average rank for the three metrics (CONSENSUS). Accordingly, 186 genes were considered as DTLs, all of which were up-regulated on syngas vs fructose (**Supplementary Table 4**). Thus, DTLs covered 42% of all the genes among the top 5% ranks of the three metrics (**Fig. 2c**). Expectedly, DTLs showed strongest enrichment for genes involved in translation (**Fig. 2e**), indicating reasonable definition of DTLs^20^.

### Transcriptional and translational control for autotrophy are uncoupled

Comparing DTRs with DTLs revealed that ∼47% of genes (166) were differentially regulated exclusively at transcript levels and ∼50% (176) exclusively at the level of translation (**Fig. 3a**). While no DTLs were down-regulated on syngas vs fructose, the concordantly up-regulated six genes included the Fd-NAD^+^ oxidoreductase Rnf complex subunit RsxC (LABRINI_16130) and the NADH-quinone oxidoreductase subunit NuoF (07815) (**Supplementary Tables 3 and 4**). This very low ∼2% overlap clearly shows that transcriptional and translational regulation involved in realising autotrophy in *C. autoethanogenum* are uncoupled. Consistently, no enriched functional categories are shared between DTRs and DTLs (**Fig. 3b**). Furthermore, the enrichment data show that two key metabolic pathways for autotrophy – acetate & ethanol production and energy conservation – are only regulated at transcript levels while ribosomal proteins are only regulated at translational level (**Fig. 3b**). The former includes various alcohol and aldehyde dehydrogenases (including Adh4 (09125), the most abundant Adh enzyme in *C. autoethanogenum*)^13^, the bifunctional AdhE1 (18700), and the aldehyde Fd oxidoreductase AOR2 (00495). Transcriptional regulation of these genes was also seen with faster autotrophic growth of *C. autoethanogenum*^33^. Additionally, strictly transcriptional regulation was used to up-regulate components of the Rnf complex (16135⍰16155), that generates proton motive force to drive the ATPase in *C. autoethanogenum*^63,64^ (see also **Fig. 6**). Transcripts of the Rnf complex are also up-regulated in *C. autoethanogenum* syngas vs fructose batch cultures^55^. Interestingly, among the translationally regulated ribosomal proteins, S7 (09680) is known to act as an autogenous translation repressor^65^ while L35 (06570) can act in an extra-ribosomal context to inhibit ornithine decarboxylases^66^, which support acid-resistance and maintenance of optimal growth in Clostridia^67^. Additionally, L7Ae (09690) participates in tRNA processing, RNA modification, and translational regulation in archaea^68^. Only four genes showed discordant changes between transcriptional and translational regulation, with 07435 and 00130 putatively involved in iron metabolism^69,70^, suggesting potential buffering effects (**Supplementary Tables 3 and 4**).

**Fig. 3.**
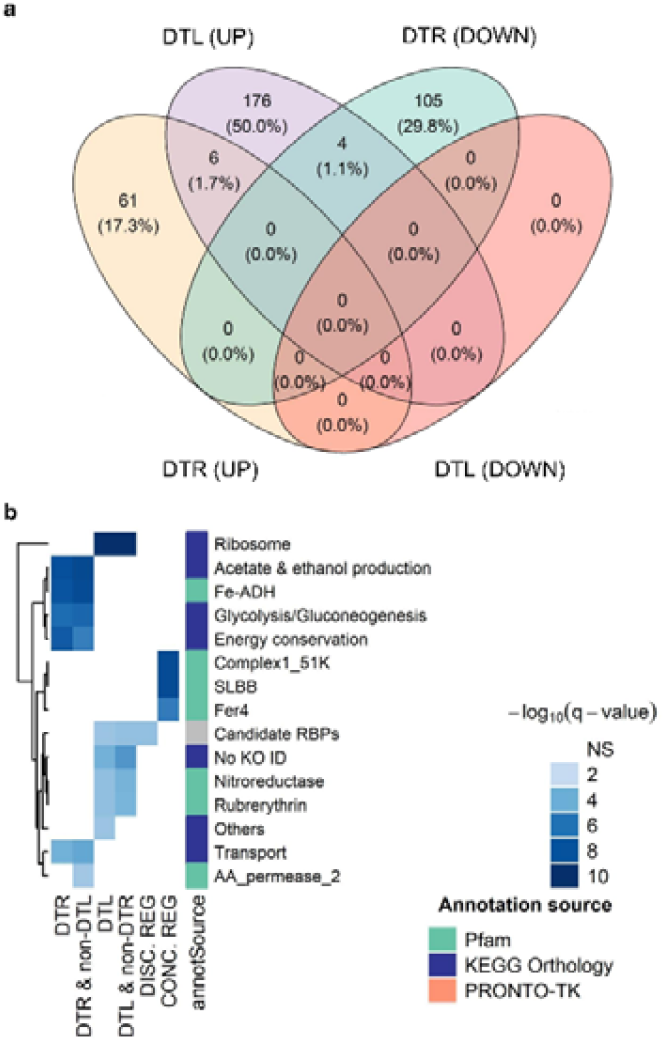
Uncoupled transcriptional and translational control for autotrophy of *C. autoethanogenum*. **a** Venn diagram showing overlaps between up- or down-regulated differentially transcribed genes (DTR) and differentially translated genes (DTL) between syngas and fructose chemostats. b Functional classes enriched in DTRs, DTLs, in genes exclusively regulated at transcriptional (DTR & non-DTL) or translational (DTL & non-DTR) level, and in genes which were both DTRs and DTLs showing concordant or discordant changes by hypergeometric test (FDR = 5%). Pfam^61^, Pfam entries from eggNOG-mapper 2.1.12^62^ ; KEGG Orthology, KEGG Orthology (KO) functional categories^60^ from Valgepea *et al*. (Level 3 in Table S3^13^ ); NS, not significant. PRONTO-TK, computationally predicted RNA-binding proteins by PRONTO-TK^50^ .

**Fig. 4.**
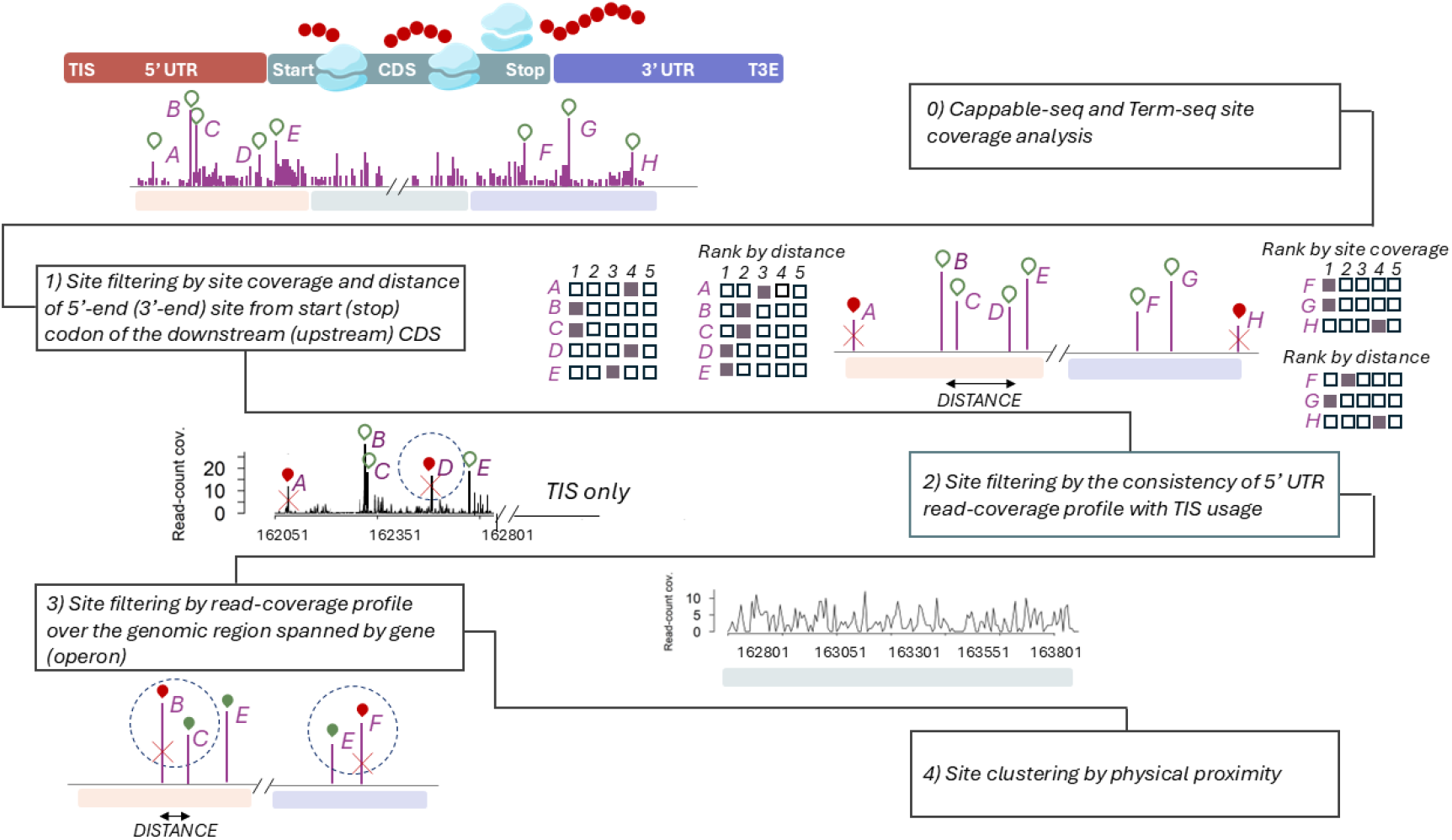
Heuristic procedure used in this work for genome-wide identification of transcription initiation (TIS) and transcription 3’ end (T3E) sites. The procedure consists of multiple filtering steps to ensure reliable identification of TISs and T3Es from preliminary sites extracted from Cappable-seq and Term-seq analysis (see Methods for details). Shortly: Filter 1) selects preliminary sites based on two criteria: site coverage in read counts and distance from the start (stop) codon of the downstream (upstream) coding DNA sequence (CDS). Sites are partitioned into five quantiles according to each criterion and quantiles for each TIS or T3E are combined. A TIS (T3E) passes to next step if the composite quantile accounting for both criteria is sufficiently high; Filter 2) is exclusively applied to 5’-untranslated regions (UTRs) and checks if it contains a stretch of sequence with a length of ≥50% of the total UTR length with a median read coverage ≥5 and ≥50% higher than for 100 nt upstream of the TIS; Filter 3) assesses the genomic region spanned by the gene (operon) for uniformity in read coverage; Filter 4) selects the TIS (T3E) closest to the start (stop) codon of the downstream (upstream) CDS for genes with multiple TISs or T3Es that are closer than 5 nt from each other.

### Mapping genome-wide transcriptional architecture

Experimental mapping of transcriptional architecture is essential to accurately identify key RNA sequences – 5’UTR and 3’UTR – through which RBPs typically facilitate post-transcriptional regulation^44,45^. We thus applied Cappable-seq^46^ to determine TISs and Term-seq^49^ to profile T3Es at genome-scale (**Fig. 1**). Sequencing metrics and read statistics with mappings are in **Supplementary Table 5**. To accurately map TISs and T3Es as primary or internal at the level of transcriptional units^71^, we organised *C. autoethanogenum* protein-coding genes into monocistronic operons with a single gene and polycistronic operons operationally defined as sets of protein-coding genes consecutively encoded on the same strand and separated by ≤60 nt. We identified 1,721 single genes while 2221 genes were organised into 781 polycistronic operons (**Supplementary Table 6**).

We next employed a rigorous and heuristic multi-step procedure to determine TISs and T3Es for each operon (**Fig. 4**; see Methods for details). Shortly, we first extracted candidate sites upstream and downstream of an operon by removing sites with close-to-background read coverage and then used a series of established criteria – distance from CDS (filter #1 on **Fig. 4**), site coverage (#1), consistency of read coverage for TISs (#2), and uniformity of read coverage across the CDS (#3) – for filtering candidate sites (and clustering if needed; #4) to assign a single primary or multiple (primary and alternative) TISs and T3Es.

Analysis of Cappable-seq data identified 1,179 unique TISs corresponding to 1,588 genes (743 monocistronic and 300 polycistronic operons) that accounts for ∼40% of protein-coding genes in *C. autoethanogenum* (**Supplementary Table 7**). From these, 1,003 were identified from syngas and 991 from fructose with 815 from both cultures. In total, around 88% TISs are primary and ∼97% map to 5’UTRs (**Fig. 5a**) while ∼90% of genes were assigned with a single TIS (**Fig. 5b**). Our results show a median 5’UTR length of 54 nt with 75% being <100 nt (**Fig. 5c; Supplementary Fig. 5a**) similar to values seen for *C. autoethanogenum* batch cultures using dRNA-seq^72^. We took advantage of the experimental TIS data for promoter motif identification using MEME^73^ for primary TISs. This supported the quality of our TIS data as we detected the well-described Pribnow box (TATAAT)^74^ motif 10 nucleotides upstream of TISs for ∼74% TISs and, at a lesser extent, the −35 motif (TTGTCA) for ∼8% TISs (**Fig. 5d**). Notably, the TG dinucleotide in the Pribnow box^75,76^ and the AT-rich region downstream of the −35 motif^77^ stimulate transcription.

**Fig. 5.**
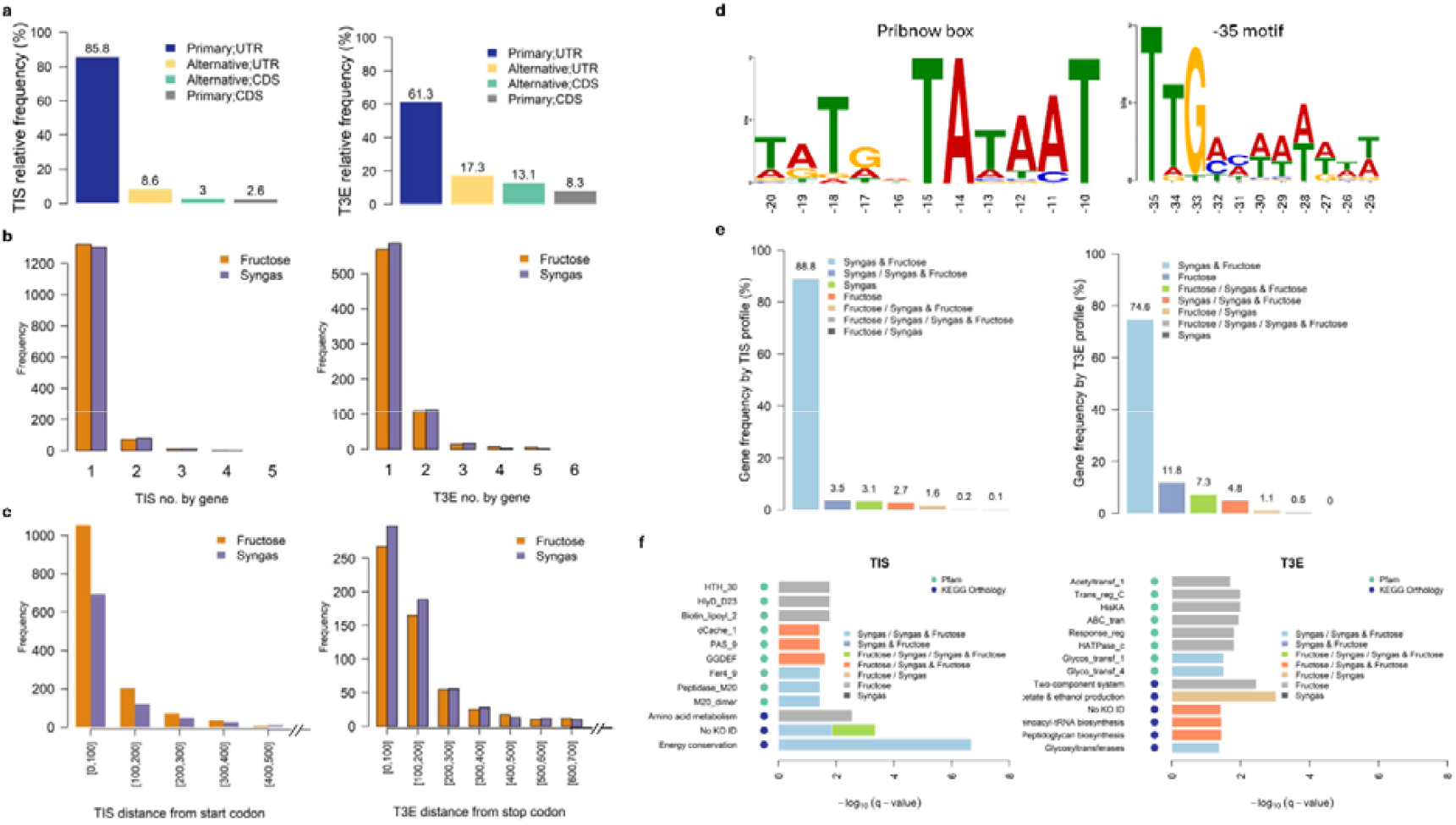
Characteristics of transcriptional architecture in *C. autoethanogenum*. **a** Classification of transcription initiation (TIS) and transcription 3’ end (T3E) sites according to strength (primary or alternative) and localization (UTR or coding DNA sequence, CDS). Primary site shows highest site coverage and shortest distance from CDS start (stop) codon while remaining sites passing the filtering procedure shown on Fig. 4 are identified as alternative sites. **b** Frequency of TISs and T3Es per gene for syngas and fructose chemostats. **c** Distribution of 5’ and 3’UTR lengths for syngas and fructose chemostats (5’UTR < 500 nt; 3’UTR < 700 nt; see **Supplementary Fig. 5** for full distribution). **d** Promoter motifs identified in 5’UTRs by MEME^73^. **e** Frequency of gene-specific TIS or T3E profiles for syngas and fructose chemostats. Bars are aggregate of merged data from biological triplicate chemostat cultures. Syngas & Fructose, site(s) shared between two conditions; Syngas / Syngas & Fructose, site(s) unique to syngas and site(s) shared between two conditions; Syngas, site(s) unique to syngas; Fructose, site(s) unique to fructose; Fructose / Syngas & Fructose, site(s) unique to fructose and site(s) shared between two conditions; Fructose / Syngas / Syngas & Fructose, site(s) unique to syngas, site(s) unique to syngas, and site(s) shared between two conditions; Fructose / Syngas, site(s) unique to syngas and site(s) unique to syngas. **f** Functional classes enriched in genes defined by TIS and T3E profile by hypergeometric test (FDR=5%). Pfam, Pfam^61^ entries from eggNOG-mapper 2.1.12^62^ ; KEGG Orthology, KEGG Orthology (KO) functional categories^60^ from Valgepea *et al*. (Level 3 in Table S3^13^ ).

Analysis of Term-Seq data identified 751 unique TE3s, 620 from syngas and 621 from fructose with 490 from both cultures, for a total of 765 genes (**Supplementary Table 7**). Overall, ∼70% T3Es are primary (**Fig. 5a**), ∼71% of genes were assigned with a single T3E (∼23% with two) (**Fig. 5b**), and the median 3’ UTR length was 122 nt with 75% being <241 nt (**Fig. 5c; Supplementary Fig. 5b**). Interestingly, ∼21% of T3Es mapped to CDSs (**Fig. 5a**), indicating potentially premature termination or processing^78^, that was dependent on the growth substrate for 46 monocistronic and 15 polycistronic operons functional in transcription, chemotaxis, transport, amino acid metabolism, and glycolysis/gluconeogenesis (**Supplementary Table 8**). We note that, to the best of our knowledge, determination of T3Es has been performed only once before for acetogens^79^.

We show the localization of TISs and T3Es for genes in key pathways for autotrophy on **Fig. 6**. More broadly, **Supplementary Table 9** contains coordinates and annotations of CDSs, TISs, and T3Es to allow intuitive exploration of *C. autoethanogenum* transcript boundaries by any visualisation tool, for instance with the open-source Integrative Genomics Viewer (IGV)^80^.

### Differential usage of TISs and T3Es for autotrophy and heterotrophy

To explore the extent of differential usage of TISs and T3Es between autotrophy and heterotrophy, we next classified genes according to the growth substrate specificity of their TISs and T3Es. The large majority of genes (∼89%) feature a single TIS used for both syngas and fructose (**Fig. 5e**; light blue) while only a ∼6% minority of genes use TISs exclusive to syngas (∼3%; green) or fructose (∼3%; orange), which includes a transcription elongation factor (07115) (**Supplementary Table 10**). The remaining set of genes show a composite profile of TIS usage with both TISs exclusive and common for either substrate. Similarly to TIS data, most genes (∼75%) feature a single T3E for both syngas and fructose (**Fig. 5e**; light blue) (**Supplementary Table 11**). However, RNA processing at 3’UTRs appears to be more dependent on growth substrates as ∼12% of genes show TISs exclusive to fructose (light purple) with another ∼7% which show exclusive TISs for fructose along with shared TISs (green). Interestingly, a glyceraldehyde-3-phosphate dehydrogenase (08720) with an accompanying transcriptional regulator (08725) showed the most versatile T3E usage with both T3Es exclusive and common for either substrate (gray). Notably, we did not detect a single gene showing only T3Es exclusive for syngas.

Importantly, we detected enrichment of genes involved in energy conservation in the group of genes featuring syngas-specific TISs in addition to TISs shared for both substrates (**Fig. 5f**). This includes genes encoding the Rnf complex, a key activity for autotrophy in acetogens^63,64^ (**Fig. 6**). Furthermore, genes with syngas-only TISs include several genes with Pfam domains involved in two-component signal transduction systems (TCS) as they encode transcriptional regulators (domain ‘Trans_reg_C’), histidine kinases (HK; ‘HisKA’), and response regulator (RR; ‘Response_reg’) proteins. Autophosphorylation of a HK by signal sensing drives the phosphorylation and activation of a RR that functions as a molecular switch to control diverse adaptive responses^81,82^. Notably, gene with ID 04855 harbours the ligand-binding domains GAF and PAS, which can bind carbon monoxide^83^, and the HK-A domain, which is the phospho-acceptor domain of HKs. The two genes upstream – 04850 and 04855 – both contain the REC domain, which receives the signal from a HK. An additional TCS was enriched here with 16335 containing a REC domain and 16340 containing the HK-A domain. These syngas-only TISs could potentially be controlled by the σ^54^-dependent Fis family transcriptional regulator (02225) since it also has a syngas-only TIS and a σ^54^ interaction domain occurs repeatedly in TCS genes^84,85^. These are potentially relevant observations to understand transcriptional regulation of autotrophy in acetogens as we previously isolated *C. autoethanogenum* strains with improved phenotypes from autotrophic adaptive laboratory evolution carrying mutations in HKs and RRs^35^, which might be part of yet-to-be-described regulatory networks^86^.

Only two genes displayed both a syngas-specific and a fructose-specific TIS: the radical SAM protein (11570) and a transcription elongation factor (07115), the latter being consistent with the identification of transcriptional regulators (02225, 12490, 13845, 02980, 18605, 18630) which are associated with substrate-specific TISs. Notably, genes featuring exclusively fructose-specific T3Es were enriched with TCSs, some of which include genes encoding HAMP-domain containing HKs (12620-12625, 03770-03775), which are putatively involved in the coordination of cell wall remodeling during cell division in Gram-positive bacteria^87,88^. For T3Es, the functional category ‘acetate & ethanol production’ was enriched in genes displaying TE3s specific to either syngas or fructose (**Fig. 5f**).

## Transcription-initiation signals for the Rnf complex are enhanced for autotrophy

Our analysis detected TISs and T3Es for genes in key metabolic pathways and activities for autotrophic growth of *C. autoethanogenum*: acetate and ethanol production, energy conservation, the WLP, and hydrogenases (**Fig. 6**). Comparison of the strength of transcription-initiation and termination signals between the two substrates could reveal regulation critical for autotrophy.

Among the TISs identified in both syngas and fructose, we detected significant differences (q < 0.05) in the expression levels of 155 TISs linked to 144 genes between the two substrates (**Supplementary Table 7**). Strikingly, transcription-initiation signals for components of the Rnf complex, which plays a key role in energy conservation for autotrophy^64,89,90^, were strongly enhanced on syngas vs fructose (**Fig. 6; Supplementary Fig. 6a**). Thus, the stronger initiation signals most likely lead to the enhanced expression of the Rnf complex CDS transcripts (**Fig. 3B; Supplementary Table 3**). Interestingly, genes with weaker initiation signals on syngas were enriched with CooS1 carbon monoxide dehydrogenase subunits (15005⍰15015), genes involved in ethanol production (02655, 16430, 18700, and 19700), genes belonging to glycolysis/gluconeogenesis, and numerous transcription factors (**Fig. 6; Supplementary Fig. 6a**). Importantly, overall, the change in transcription-initiation signal strength is consistent with differential expression of respective transcripts, as shown by the enrichment of genes with differential TIS signals among the top or bottom ranks of transcript expression changes between the two substrates (**Supplementary Fig. 7**).

**Fig. 6.**
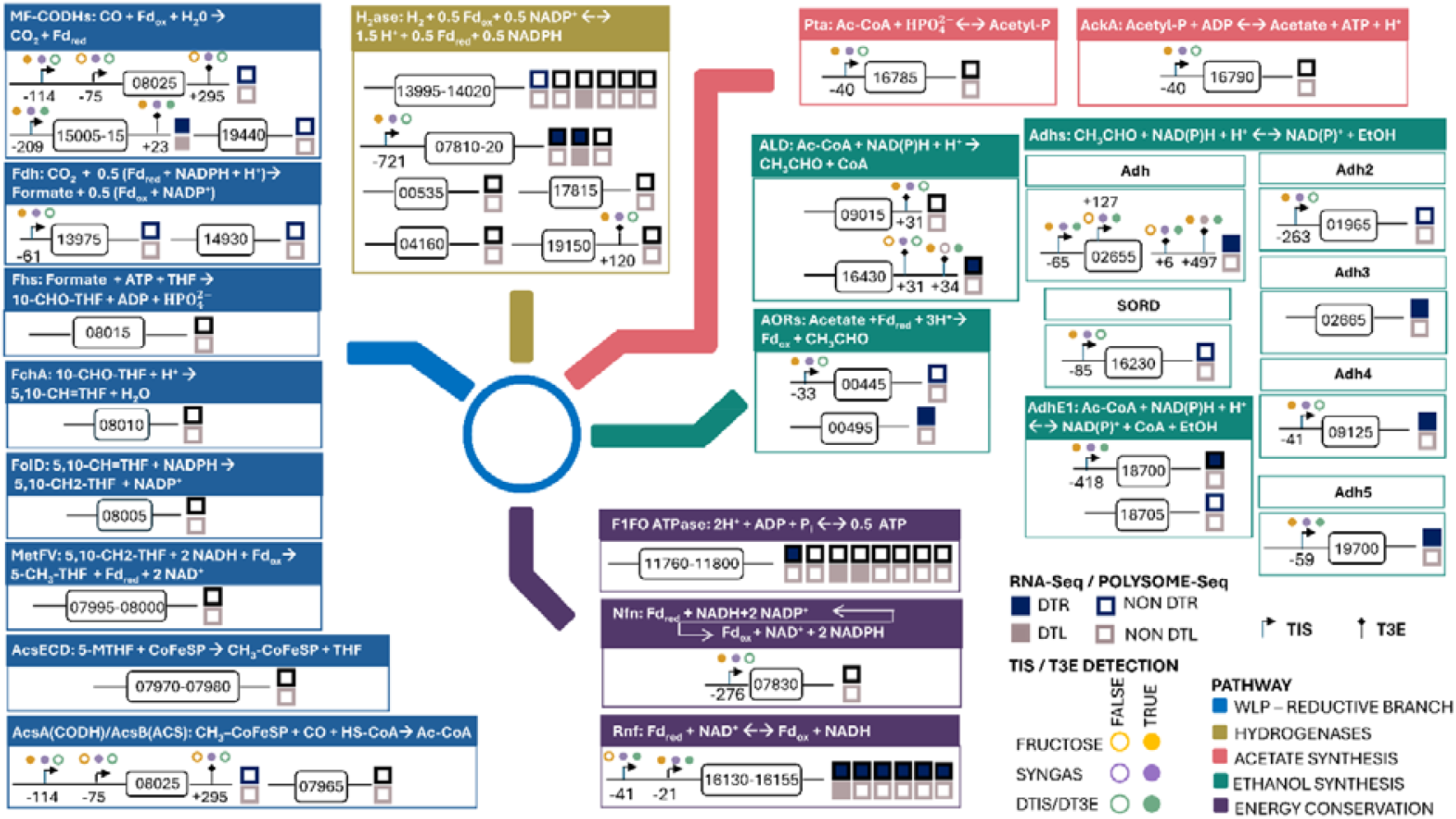
Transcriptional architecture and transcriptional/translational regulation in key metabolic pathways for autotrophy of *C. autoethanogenum*. Boxed numbers are LAbrini gene IDs. Transcription initiation (TIS) or transcription 3’ end (T3E) site distance shown from start codon of downstream coding DNA sequence (CDS) or stop codon of upstream CDS, respectively. DTIS, differentially expressed TIS; DT3E, differentially expressed T3E; MF-CODH, monofunctional carbon monoxide dehydrogenase; Fd, ferredoxin; 10-CHO-THF, 10-formyl tetrahydrofolate; 5,10-CH=THF, 10-methenyl-THF; 5,10-CH2-THF, 10-methylene-THF; 5-CH_3_-TFH, 5-methyl-THF; Ac-CoA, acetyl-CoA; EtOH, ethanol. See **Supplementary Table 1** for gene descriptions.

**Fig. 7.**
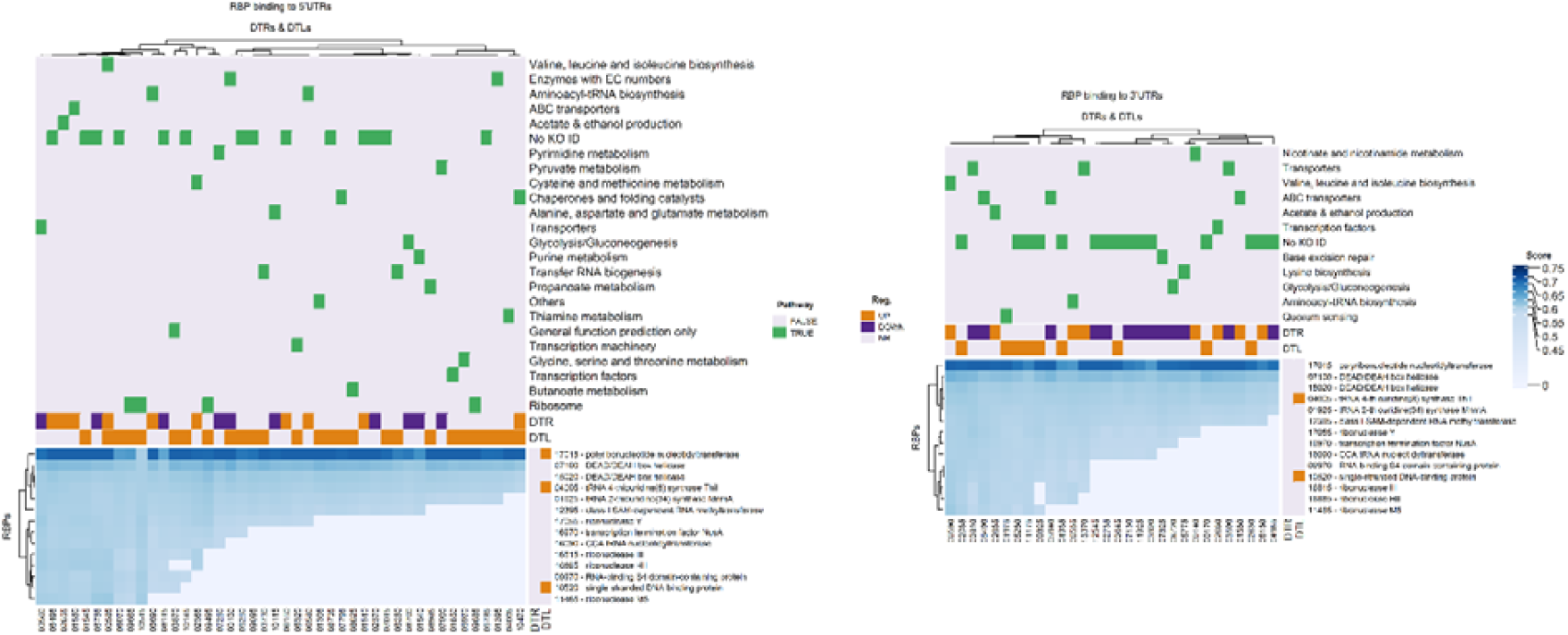
RNA-binding proteins (RBPs) contributing to autotrophy of *C. autoethanogenum*. Predicted RBP interactions with 5’ and 3’UTRs of differentially transcribed genes (DTRs) and differentially translated genes (DTLs) with RBPs in rows and target genes in columns. Blue heatmap shows catRAPID scores for RBP-RNA interactions. Row dendrogram on the left side and column dendrogram on the top of the heatmaps are from complete-linkage hierarchical clustering of the Euclidean distance matrix based on RBP-RNA interaction scores computed by catRAPID. KEGG Orthology functional categories^60^ from Valgepea *et al*. **(Level 3 in Table S3**^13^) for target genes are shown on top. See **Supplementary Table 1** for target gene descriptions.

Within the T3Es detected in both syngas and fructose, we measured significant differences in the expression levels of 439 T3Es linked to 341 genes between the two substrates (**Supplementary Table 12**). Genes with enhanced termination signals for autotrophy included a gene cluster for cobalamin biosynthesis (05425⍰05445) while genes with repressed termination signals were enriched with ABC transporters, flagellar proteins, and genes involved in cell wall metabolism (**Supplementary Fig. 6b**). The relevance of these changes in termination signal strengths for autotrophy remain unclear.

### RNA-binding proteins (RBPs) involved in transcriptional and translational regulation for autotrophy

Accurate mapping of transcriptional architecture defines UTR RNA sequences where RBPs could bind to execute post-transcriptional regulation. We could identify 5’UTRs for ∼66% of DTRs and ∼61% of DTLs, and 3’UTRs for ∼28% of DTRs and ∼10% of DTLs. To identify potential RBP binding regions in UTRs and thus RBP-RNA interactions, we first predicted potential RBPs for *C. autoethanogenum* using PRONTO-TK^50^, a neural network-based protein function prediction tool (**Fig. 1; Supplementary Table 13**). PRONTO-TK uses ProtT5-XL protein language model embeddings to encode sequence-derived biochemical and structural features. We trained a binary classifier using the RNA binding GO term and its descendants, based on experimentally validated protein annotations from the UniProt knowledgebase^91^. We then used the computational tool catRAPID^51^ to search for candidate RBP binding sites in the UTRs of DTRs and DTLs (**Fig. 1; Supplementary Table 14**), while excluding structural ribosomal proteins from the RBP list. Overall, 5’UTRs of ∼11% of DTRs (20 genes) and ∼15% of DTLs (27 genes) were predicted to contain RBP binding sites while the values for 3’UTRs were 12% for DTRs (21) and ∼5% for DTLs (9) (**Fig. 7; Supplementary Table 14**). Most RBP target genes were regulated exclusively transcriptionally (34 out of which 13 up-regulated and 21 down-regulated on syngas vs fructose) or translationally (32 up-regulated) **(Fig. 7**). The remaining three RBP targets were a ferritin-encoding gene (00130) showing different regulation between transcription and translation, and a heat-shock protein (10470) and a ketol-acid reductoisomerase (00585) showing concordant regulation. Interestingly, 00585 is among the top 10 most abundant proteins in *C. autoethanogenum*^13^. Most RBPs recognised cis-regulatory elements in both 5’ and 3’UTRs of target genes while individual genes were rarely bound by RBPs in both of their UTRs. Within the latter, a potentially relevant example for autotrophy is the alcohol dehydrogenase 02655. While genes involved in coenzyme synthesis for redox reactions, quorum sensing, and base excision repair were mediated by RBPs exclusively via 3’UTRs, translation-related genes (e.g. amino acid metabolism, tRNA processing) and transporters showed binding events in both UTRs.

The above-mentioned UTRs showed binding sites for 14 RBPs while none were found to bind uniquely either to 5’UTRs or 3’UTRs (**Fig. 7**). Furthermore, none of the 69 target genes were linked to regulation by a single RBP. Notably, three RBPs were among the translationally up-regulated DTLs on syngas: the polynucleotide phosphorylase PNPase (17015) involved in mRNA turnover^92^ and RNA-mediated regulation^93^; a protein annotated as a single-stranded DNA-binding protein (10520); and the 4-thiouridine synthetase ThiI (04005), which stabilises tRNA folding by realizing post-transcriptional modifications of uracil at position 8^94^ and is involved in the biosynthesis of sulfur-containing biomolecules such as thiamin^95^.The remaining RBPs included proteins well-known for their roles in RNA metabolism: the S-adenosyl-L-methionine dependent RNA methyltransferase (MTase) (12395)^96^; the ribonucleases (RNases) Y (17055), III (16815), HII (16885), and M5 (11465) involved in RNA decay^97-99^; the transcription attenuation protein NusA (16970)^100,101^; the CCA tRNA nucleotidyltransferase (16090), which catalyzes the addition of the cytosine-cytosine-adenine (CCA) tail to the tRNA species that is indispensable for amino acid charging and tRNA functionality^102^; and two DEAD/DEAH box helicases (07100 and 15020) involved in RNA decay^103^.

None of the RBPs were predicted to bind to genes involved in the following key activities for autotrophy of acetogens: C_1_-fixation using the WLP, energy conservation through the Rnf complex, redox homeostasis through the Nfn transhydrogenase, or the various hydrogenases. At the same time, multiple RBPs showed UTR interactions with three alcohol dehydrogenases (08985, 08625, and 02655) (**Fig. 6**) that can be critical for maintenance of redox and energy homeostasis for *C. autoethanogenum* autotrophy through regulation of carbon flux between acetate and ethanol^14,104^, the two major metabolic by-products (**Supplementary Fig.1**). Expectedly, several RBPs targeted translation-related processes, including ribosomal structural proteins, tRNA synthetases, genes involved in amino acid metabolism and in biosynthesis of tRNA hypermodified nucleosides, namely queuosine and *N*^6^-threonylcarbamoyladenosine, which are essential for modulating protein translation^105^ (**Fig. 7**). Relevant to autotrophic biomass formation through gluconeogenesis was the binding of several RBPs to the 3’UTR of the type I glyceraldehyde-3-phosphate dehydrogenase (08720).

## Discussion

While post-transcriptional regulation of gene expression plays an important role in acetogens^60^, none of the key players in this regulatory layer – RBPs – had been described before. We thus also lacked knowledge about RBP-RNA interactions in acetogens that could facilitate global adjustments in metabolism, for instance towards autotrophy. To address these gaps with this work, we employed a comprehensive approach by combining chemostat cultures, functional genomics, and computational methods to identify genome-wide RBPs and RBP-RNA interactions contributing to autotrophy in the model-acetogen *C. autoethanogenum*.

We focused our approach on genes displaying transcriptional and/or translational regulation by their transcripts being differentially transcribed and/or translated during autotrophy vs heterotrophy. Widespread translational regulation could imply its dominant role for regulating protein levels for realising autotrophy. We determined a low number of DTLs (186), which is consistent with the good correlation (R ∼ 0.69 for 1,797 genes) between transcript levels quantified in syngas chemostats here and protein levels in LAbrini syngas chemostats in previous works^34,86^ (**Supplementary Fig. 8**), and with the low number of proteins differentially expressed between CO, CO+H_2_, and CO+CO_2_+H_2_ grown *C. autoethanogenum* chemostats^106^. Indeed, metabolic flux adjustments between the latter cultures are dominantly accompanied by changing enzyme catalytic rates rather than concentrations^13^. Such post-translational regulation of fluxes is also seen in self-oscillating *C. autoethanogenum* autotrophic cultures^12,14^ and this is presumably the most efficient regulatory mode of fluxes for energy-limited metabolism since close to half of ATP for biomass proliferation is used for protein synthesis^107,108^ . Furthermore, post-translational regulation would also allow cells to more rapidly react to gaseous substrates becoming available in natural environments by keeping a higher baseline expression of key proteins. At the same time, nearly half of transcripts showed statistically different expression (q-value < 0.05) here (**Fig. 2a; Supplementary Table 3**), consistent with data from *C. autoethanogenum* syngas and fructose batch cultures^55^. Thus, transcriptional regulation seems to be required to ensure the baseline proteome expression for both autotrophy and heterotrophy. Notably, transcriptional control of Rnf complex components and alcohol and aldehyde dehydrogenases detected in our work (**Fig. 6**) suggests attractive strain engineering targets in key metabolic activities of *C. autoethanogenum*.

Overall, ∼19% of differentially regulated genes showed interactions with RBPs in our dataset (**Fig. 7**). The fact that we did not identify any RBPs binding only to genes with key activities for autotrophy (e.g. WLP, Rnf, Nfn, hydrogenases) implies that acetogens, at least *C. autoethanogenum*, do not seem to use an RBP-level mechanism for specifically regulating expression of genes exclusively needed for autotrophy. Although multiple RBPs targeted alcohol dehydrogenases, potentially critical for maintenance of redox and energy homeostasis^14,104^, elucidation of key RBPs here, however, would be highly challenging due to the large number of candidate RBPs, their binding to other target genes with potentially essential functions, and the difficulties with creating such a number of *C. autoethanogenum* gene deletion mutants. Importantly, our study captured several RBP-RNA interactions well-known in bacteria that are needed for optimal growth in general. These interactions are therefore also involved in mediating differential gene transcription and translation profiles for realising autotrophy in *C. autoethanogenum* and will be thus discussed in more detail next.

Several differentially regulated genes in our dataset showed binding by the RBP NusA (**Fig. 7**; 16970), a factor for transcription attenuation where bacteria direct the RNA polymerase to either terminate transcription or transcribe downstream genes in an operon as a response to metabolic signals^100,101^. In contrast to translation-controlled termination of transcription in *E. coli*, the Gram-positive model microbe Bacillus subtilis (*C. autoethanogenum* is Gram-positive) lacks kinetic coupling between the transcription elongation factor and the pioneering ribosome^109^. Therefore, operon-specific RBPs^110-113^, or trans-acting factors, such as NusA^78,114^ are expected to support transcription attenuation. Our study shows that NusA is involved in realising autotrophy in *C. autoethanogenum*, as it is predicted to bind to genes involved in metabolic processes that are sensitive to transcriptional attenuation such as biosynthesis of thiamine, amino acids, tRNA synthetases, and ribosome biogenesis^115^. Transcription termination is often triggered by the arrangement of one of the two mutually exclusive stem structures in the nascent transcript, where the second stem is followed by a poly-U sequence forming a terminator and the second strand of the first stem base-pairs with the first strand of the second stem^116^. Indeed, we detected the presence of this characteristic arrangement in the 5’UTR of three genes targeted by NusA in our study using PASIFIC^117^: 06495, 01550, and 02365 encoding a tyrosine-tRNA ligase, a transmembrane protein, and a homoserine O-succinyltransferase, respectively. A recent study suggests that NusA-mediated stabilization of transcriptional pausing and attenuation could be involved in the establishment of transcription-translation coupling through the interaction between NusA and translation initiation factor IF2^118^. This mechanism is unlikely in *C. autoethanogenum* as B. subtilis lacks coupling between transcription and translation^109^. Thus, further studies are required to elucidate the role of NusA in *C. autoethanogenum*.

Our results further show binding of an array of RNases to differentially regulated genes (**Fig. 7**). RNases are essential mediators in RNA maturation, quality control, and degradation where changes in tRNA and rRNA maturation impact translational regulation while changes in mRNA stability affect transcript levels. mRNA decay in Gram-positive bacteria relies on the cooperation between endoribonucleases, which cleave mRNAs and provide access to 5’ and 3’ extremities, and 5’→3’ and 3’→5’ exoribonucleases, which degrade the cleaved RNA fragments^26,119^. Our data for *C. autoethanogenum* is consistent with this model as we identified binding of both endoribonucleases (RNase Y, 17055; RNase III, 16815; and RNase M5, 11465) and exoribonucleases (PNPase, 17015 and RNase HII, 16885).

The most target genes were detected for RNase Y, the major decay-initiating endonuclease in *B. subtilis* and many Firmicutes^99,120^. Supporting proteins are proposed to be important for RNase Y target selection in the Gram-positive *Staphylococcus aureus*^121^ since RNase Y appears to be strongly influenced by features distal to the position of cleavage, possibly facilitated by RNA secondary structure and RNA-protein interactions^122^. Our study suggests that this could be the case for *C. autoethanogenum* as well since RNase Y target genes were bound also by RNA helicases and RNA methyltransferases (**Fig. 7; Supplementary Table 14**). Interestingly, while RNase M5 in B. subtilis is thought to be almost exclusively active in 5S rRNA processing^123^, we detected its binding to an alcohol dehydrogenase, multiple transporters, S6 and S10 ribosomal proteins, and genes in folate and thiamine metabolism (**Fig. 7; Supplementary Table 14**). Notably,

∼88% of RNase M5 targets were also targeted by RNase III. While RNase III was initially assigned a secondary role in *B. subtilis*^124^, it has now been shown to be important in *B. subtilis* for post-transcriptional catalytic and binding activities^98^ and to be involved in translational regulation in *S. aureus*^125^. Thus, it is likely that all the three described endoribonucleases are relevant for autotrophy of *C. autoethanogenum*. For degradation of the cleaved RNA, *C. autoethanogenum* seems to primarily rely on the exoribonuclease PNPase as it targeted all the differentially regulated genes that the three endoribonucleases targeted (**Fig. 7**). What is more, PNPase was translationally up-regulated on syngas (**Fig. 7**). Use of the phosphorolytic enzyme PNPase would be beneficial for an energy-limited metabolism as the PNPase degradation products nucleoside diphosphates conserve energy^26^. Interestingly, PNPase can also play a role in complex cell adaptive responses through its recruitment in different multiprotein complexes^92^. RNA degradation could also involve the poorly characterised exoribonuclease RNase HII as it shared an unexpectedly high fraction of targets in common with RNase Y (89%), RNase M5 (67%), and RNase III (90%).

In terms of post-transcriptional RNA modifications for differentially regulated genes, the RBP pseudouridine synthase (09970) could be relevant (**Fig. 7**) as pseudouridine is a ubiquitous bacterial RNA modification, primarily on tRNA and rRNA, and secondarily on mRNA^126^. Whereas the modification plays a role in mammalian transcript stabilization^127^ and modulation of transcript translatability^128,129^, its potential effects on mRNA function in bacteria are largely unexplored^130^. The fact that 09970 targeted both DTRs and DTLs in our dataset suggests that pseudouridine can affect both transcription and translation in bacteria, similarly to mammals. Its scope seems wide for autotrophy as 09970 targets included an alcohol dehydrogenase, transporters, the threonine-tRNA ligase, and the 30S ribosomal protein S10, which is part of the processive rRNA transcription and antitermination complex.

We believe our work to be important for two reasons. Firstly, we provide the first systems-level steady-state datasets for an acetogen comparing heterotrophic and autotrophic growth. Importantly, this combined description of transcriptome expression, polysome profiles, and transcriptional architecture advances fundamental understanding of gene function, gene expression regulation, and metabolism of the ancient metabolism of acetogens. Secondly, we report, for the first time for acetogens, genome-wide protein-RNA interactions that contribute to autotrophy in a model acetogen. This is important as it provides novel targets for metabolic engineering of acetogen cell factories from the unexplored space of RBPs. Future work will need to experimentally quantify the effects of genetic perturbations of candidate RBPs on acetogen autotrophy, including potentially synergistic or epistatic links between RBPs.

## Methods

### Bacterial strain and cultivation conditions

The *C. autoethanogenum* strain LAbrini^35^ deposited in the German Collection of Microorganisms and Cell Cultures (DSM 115981) was used in this work.

*C. autoethanogenum* was grown in biological triplicate chemostat continuous cultures either autotrophically on syngas (50% CO, 20% H_2_, 20% CO_2_, and 10% Ar; AS Linde Gas) or heterotrophically on fructose (15 g/L) in a chemically defined medium (without yeast extract) as described before^104^ with the following difference for fructose cultures: CO_2_ sparged at 50 mL/min for 20h post-inoculation followed by sparging N_2_ at 50 mL/min for the rest of the fermentation. Shortly, all cultures were grown under strictly anaerobic conditions at 37°C and at pH 5 in 1.4 L Multifors bioreactors (Infors AG) at a working volume of ∼750 mL connected to a Hiden HPR-20-QIC mass spectrometer (Hiden Analytical) for online off-gas analysis. We used a syngas flow rate of 50 mL/min and agitation of 675 RPM for autotrophic and 15 g/L of fructose for heterotrophic cultures (agitation of 200 RPM) to achieve steady-state biomass concentrations of 1.37 ± 0.003 and 1.03 ± 0.008 gram of dry cell weight per litre for syngas and fructose cultures, respectively, for chemostats at dilution rates (D) 1.03 ± 0.05 and 0.997 ± 0.003 day^-1^ for syngas and fructose cultures, respectively. Steady-state results and samples were collected after culture optical density, gas uptake, and production rates had been stable for at least three working volumes.

### Cappable-seq

*C. autoethanogenum* chemostat cultures were sampled for Cappable-seq by pelleting either 5 (syngas cultures) or 6.5 mL (fructose cultures) of culture broth (to match biomass amounts) by centrifugation (5,000 × g for 3 min at 4°C), discarding the supernatant, and flash-freezing in liquid N_2_ followed by storage at ⍰80°C until analysis. Total RNA from frozen samples was extracted by using the RNeasy Mini Kit (Qiagen) according to the manufacturer’s instructions with in-column DNase treatment with the RNase-Free DNase Set (Qiagen). Total RNA samples were then converted to Cappable-seq libraries as described before^46^. Following enrichment and decapping, RNA samples were converted to Illumina-ready libraries using the SMARTer smRNA-seq kit (TakaraBio). Library qualities were assessed by the Agilent 2100 Bioanalyzer and finally libraries were sequenced using the Illumina Novaseq 6000 sequencer with 1 x 100 bp cycles (single-end).

### Term-seq

The same cell pellets as for Cappable-seq were used for Term-seq (see above for sampling details). Ribosomal RNA was depleted from total RNA using the Illumina RIBO-Zero plus rRNA depletion kit, following manufacturer instructions. Ribosomal RNA depleted-RNA 3’ ends were polyadenylated, followed by RNA fragmentation, end-repair, and library preparation using the SMARTer smRNA-seq kit (TakaraBio). Library qualities were assessed by the Agilent 2100 Bioanalyzer and finally libraries were sequenced using the Illumina Novaseq 6000 sequencer with 2 x 100 bp cycles (paired-end).

### Data analysis for genome-wide identification of TIS and T3E sites

Firstly, Term-seq reads (R1 or R2) were all trimmed of sequence elements added during library construction. Of each R1/R2 pair, trimmed-selected reads include either the R1 read, whenever the R1 contained the 3’ adapter (i.e., when the read reaches the end of the mRNA fragment), or the inverse-complement of the R2 read, which always includes the end of the mRNA fragment. Whenever possible, R1 was preferred over R2 because R1 generally has higher sequence quality. Trimmed Cappable-seq reads (by the position of their 5’-end) and trimmed-selected Term-seq reads (by 3’-terminal position) were mapped to the genome of C. autoethanogenum strain LAbrini (GenBank CP110420.1) using STAR^131^. Read quality control was carried out by Fastq and sequencing metrics and read statistics with mappings are in **Supplementary Tables 34⍰36**.

Secondly, *C. autoethanogenum* coding DNA sequences (CDS) were organized into monocistronic operons with a single gene and polycistronic operons operationally defined as sets of protein-coding genes consecutively encoded on the same strand and separated by ≤60 nt. For the analysis of Cappable-seq read coverage relative to operons, we implemented the following heuristic procedure:

1. For each operon, we considered genome regions comprising the location of the gene operon and its 5’ flanking (upstream) sequence up to the upstream gene operon on the same strand. Based on the genome annotation, we characterized each position within this region as hypothetical ‘5’-UTR’, coding region (‘CDS’), or intergenic region (‘IG’) for polycistronic operons.
2. From the hypothetical ‘5’UTR’ region, we identified the position *i* with maximum Cappable-seq 5’-end coverage 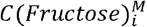 summing coverages of the three fructose bio-replicates with maximum coverage 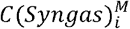 summing over the three syngas bio-replicates.
3. We identified all positions 1 within ‘5’UTR’, ‘CDS’, or ‘IG’ regions with coverages *C* (*Fructose*)_*j*_ or *C* (*Syngas*)_*j*_ such that 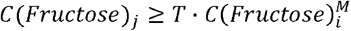 and 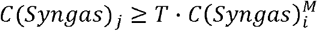, where *T* = 0.1 is a fractional threshold of the respective maximum coverages identified in the 5’UTR of the operon. We considered all sites identified in the ‘5’UTR’ from fructose or syngas bio-replicates as potential TIS candidates, and all sites identified in ‘CDS’ and ‘IG’ regions as representing background coverages.
4. We characterized each site identified within the ‘5’UTR’ region by its ranking relative to the coverages of those selected from ‘CDS’+’IG’ regions for each operon (‘*p*-value’). Thus, if 100 sites were identified as background in ‘CDS’+’IG’ regions, a ‘5’UTR’ site that had coverage higher than any of those coverages was assigned a *p*-value = 0.0. If, e.g., 10 sites from the ‘CDS’+’IG’ region had higher coverage, it was assigned a *p*-value = 0.1 (10/100).
5. Among all sites of the ‘5’UTR’ region, those with *p*-value ≤ 0.05 and with average coverage across syngas or fructose bio-replicates > 5 reads were selected.
6. To identify potential TIS candidates within ‘CDS’ or ‘IG’ regions for an operon, we selected all sites with coverages ≥50% of the maximum signal in the ‘5’UTR’ region and with average coverage across syngas or fructose bio-replicates > 20 reads.
7. All sites identified in adjacent sequence positions were collapsed into one site of ‘size’ equal to the number of positions collapsed.

An analogous approach was used to analyze Term-seq read coverage relative to operons, but here we considered for each operon, a genome region comprising the location of the operon and its 3’ flanking (downstream) region up to the downstream operon on the same strand. We characterized each position within this region as hypothetical ‘3’UTR’ region, coding region (‘CDS’) or intergenic region (‘IG’) for polycistronic operons and then followed the same procedure as Cappable-seq analysis, considering the ‘3’UTR’ region in place of the ‘5’UTR’ region.

Lastly, we used the following multi-step filtering procedure to determine TISs and T3Es for each operon from the candidate sites identified above (see also **Fig. 4**.).

1. *Site filtering by site coverage and distance from start (for TIS) or stop (for T3E) codon from CDS*. Here, candidate TISs and T3Es were filtered based on their distance from the start (for TIS) or stop (for T3E) codon of the downstream (for TIS) or upstream (for T3E) CDS *Dist* (*site, CDS start* / *stop codon*) and based on their coverage in read counts *SiteCov*. We arranged the sites in descending order based on *SiteCov* values and into ascending order based on *Dist* (*site, CDS start* / *stop codon*) and then separately partitioned into five quantiles according to their position and signal intensity. As a result, a hypothetical site *i* received the quantile *Q*(*Dist*)_*i*_ corresponding to its distance relative to the start (stop) codon of the downstream (upstream) coding sequence and the quantile corresponding to log_2_ site coverage normalized by the average coverage of all sites from all fructose samples 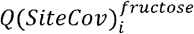 or from all syngas samples 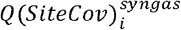. Lastly, the 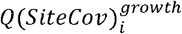 (growth is either fructose or syngas) and the *Q*(*Dist*)_*i*_ quantiles of the site *i* were summed up to define the site’s composite quantile *CQ*_*i*_ . For each gene (operon), we selected the sites displaying the lowest composite quantile that corresponds to sites which tend to display higher coverage and shorter distances from the respective start or stop codon. If the composite quantile of the site was lower than the central composite quantile, the candidate site passed to the next filtering step. We note that this procedure allows us to retain multiple and different sites per gene (operon) per substrate.
2. *Site filtering by the consistency of 5’UTR read-coverage profile with TIS usage*. Here, candidate 5’UTRs are assessed for consistency with TIS usage as 5’UTRs are expected to display non-uniform read coverage across the TISs. Thus, TISs that contained a stretch of sequence with a length of ≥50% of the total UTR length with a median read coverage ≥5 and ≥50% higher than that for 100 nt upstream of TIS passed to the next filtering steps.
3. *Site filtering by read-coverage profile over the genomic region spanned by gene (operon)*. Here, we assume read coverage over the gene (operon) to be uniform. Thus, a site passed to the next filtering steps if it is linked to a CDS with a median value of the median read-coverage values computed over 200 nt-long sliding windows being <10% different from the median read-coverage over the full CDS (except if an internal TIS (T3E) is present).
4. *Site clustering by physical proximity*. Here, we selected a representative site if multiple close sites were linked to a gene (operon). Thus, from sites closer than 5 nt from each other, the TIS (T3E) closest to the start (stop) codon of the downstream (upstream) CDS was selected as the primary site. All other sites were considered as alternative sites.

### Differential expression analysis of TIS and T3E signals

We quantified differences in TIS and T3E signals (i.e. site coverage) between syngas and fructose using EdgeR with the quasi-likelihood F-test procedure^132^. To determine sites with statistically significant differences (q-value < 0.05), we applied the Benjamini–Hochberg procedure to the F-test results, controlling the false discovery rate (FDR) correction at 5%^133^.

### Polysome-seq

#### Sample preparation and sequencing

*C. autoethanogenum* chemostat cultures were sampled, samples processed, and polysome profiling was performed as previously described^59^. Briefly, translation was arrested by adding chloramphenicol to bioreactors to a final concentration of 0.1 mg/mL. Next, 80 (syngas cultures) or 100 mL (fructose cultures) of culture broth (to match biomass amounts) in four replicate samples were harvested, washed twice, and resuspended in lysis buffer (20 mM Tris HCl pH 8, 140 mM KCl, 40 mM MgCl_2_, 0.5 mM DTT, 100 μg/mL chloramphenicol, 1 mg/mL heparin, 20 mM EGTA, 1% Triton X-100). After mechanical cell disruption with glass beads, supernatant was flash-frozen in liquid N_2_ followed by storage at ⍰80°C until analysis. Then, mRNA-ribosome complexes were size-separated on a sucrose gradient (10 to 50% w/v) in polysome gradient buffer (same composition as lysis buffer except for heparin at a final concentration of 0.5 mg/mL) into 24 subfractions. After protein denaturation and nucleic acid precipitation performed as previously described^59^, total RNA was extracted in subfractions using RNeasy midi kits (Qiagen) combined with on-column DNase I treatment. RNA amount was quantified using a Nanodrop spectrophotometer (Thermo Fisher Scientific). External ERCC RNA spikes (Invitrogen) were added to 1 µg of each total RNA sample according to manufacturer instructions. The levels of 16S and 23S rRNAs in subfractions were calculated using the Agilent 2100 Bioanalyzer 2100 and used to pool the subfractions into four fractions labelled A to D. Fraction A (from 1 to 4.5 min) consisted of subfractions containing DNA, free RNAs, and free small and large ribosomal subunits. The first peak (Fraction B from 4.5 to 6 min) was attributed to the monosome, Fraction C (from 6 to 8 min) to polysomes with low ribosome density and Fraction D (from 8 to 12 min) to polysomes with high ribosome density. The average number of ribosomes in Fractions B, C, and D was extrapolated knowing the elution time of the monosome^59^ and were found to be similar for syngas and fructose cultures (1.1 ± 0.1 ribosomes in Fraction B, 2.9 ± 0.2 ribosomes in Fraction C, and 7.7 ± 0.5 ribosomes in Fraction D). Total RNA in Fractions A, B, C, and D was ribo-depleted using the RIBO-Zero kit (Illumina). After fragmentation, the first strand cDNA was synthesized using random hexamer primers. During the second strand cDNA synthesis, dUTPs were replaced with dTTPs in the reaction buffer. The directional library was ready after end repair, A-tailing, adapter ligation, size selection, USER enzyme digestion, amplification, and purification. Finally, libraries were sequenced using the Illumina Novaseq X Plus Series platform with 2 x 150 bp cycles (paired-end).

#### Quantification of transcript abundances

Firstly, reads with >10% of undetermined bases (N) and reads with >50% of bases with Qscore ≤ 5 were excluded. Secondly, the ERCC RNA spikes were used to determine the linearity range between read counts and RNA concentration and thus features among the 4,043 LAbrini genes and 92 ERCC RNAs with <10 read counts were excluded for reliable quantification (259 and 264 features for fructose and syngas, respectively). Reads trimmed of Illumina adapters were then mapped to the genome of *C. autoethanogenum* strain LAbrini (GenBank CP110420.1) and ERCC RNA spikes using Bowtie2^134^ and read counting was performed using the featureCounts function within Rsubread^135^. To account for variations in sample preparation and analysis from ribodepletion to sequencing, two normalization steps were used. Firstly, we applied the RUV method to remove unwanted variation^136^ using the ERCC RNA spikes introduced in each fraction before the ribodepletion step: we obtained *N*_*i,j,k NormRUVg*_, with the count of the transcript *i*, in the fraction *j*, and bio-replicate *k* after normalizing with the RUV method, with *i* ⍰ {1,…,3876}, *j* ⍰ {A,B,C,D} and *k* ⍰ {1,2,3} on fructose and with *i* ⍰ {1,…,3871}, *j* ⍰ {A,B,C,D} and *k* ⍰ {1,2,3} on syngas. Secondly, we accounted for the variability in the initial amount of total RNA between fractions as a constant 0.5 µg of total RNA was taken in each fraction for ribodepletion: we obtained the final normalized count of the transcript *i*, in the fraction *j*, and bio-replicate *k* as *N*_*i,j,k*_ = *N*_*i,j,k NormRUVg*_ x total RNA *quantity*_*j,k*_ / 0.5. Finally, for each transcript *i*, we calculated the proportion of RNA copies in each fraction *j* for each bio-replicate *k*:

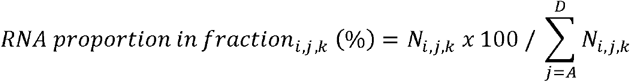

#### Consistency analysis of bio-replicates

Due to the lower-than-expected correlation between fractional mRNA counts of some bio-replicates (**Supplementary Fig. 3**), we assessed data consistency between bio-replicates within both chemostat datasets by three complementary methods suited for compositional data, such as polysome-seq: correlation analysis, compositional distance analysis, and principal component analysis (PCA). Firstly, we computed the Spearman’s rank correlation coefficient (*ρ*) between mRNA counts per polysomal gradient fractions from total counts between bio-replicates. A bio-replicate was determined as an outlier if it showed the highest number of poorly correlated polysomal fractions; a fraction was considered poorly correlated if its data correlated poorly (ρ < 0.7) with the other bio-replicates which in turn correlated between each other with ρ > 0.7. Secondly, we calculated the average of all mRNA counts for each fraction and bio-replicate, then centered-ln-ratio transformed the averages, and lastly computed the Aitchison’s distance between each pair of bio-replicates. A bio-replicate was determined as an outlier if it showed the maximal average Aitchison’s distance^137^ from the other bio-replicates that was greater than the median distance observed across syngas and fructose. Thirdly, we collated all mRNA counts for each fraction and bio-replicate as a concatenated dataset, then centered-ln-ratio transformed gene compositional data and lastly performed PCA to compute the average distances between the centroids of each pair of bio-replicates. A bio-replicate was determined as an outlier if its centroid showed the maximal average distance from the centroids of the other bio-replicates that was greater than the median distance observed across syngas and fructose. Finally, we assembled the information gathered by the three approaches, and we determined as an outlier the bio-replicate which was found to be outlier according to the highest number of approaches.

#### Determination of DTLs

We used three complementary metrics to quantify translational regulation and finally determined DTLs using a consensus approach (CONSENSUS) over the three metrics.

First, for the TL-STATUS metric, we firstly calculated for each gene its translation status in both syngas and fructose as the ratio between the sum of the mRNA proportions quantified in the ribosome-bound and ribosome-free fractions. Secondly, subtracting the fructose value from syngas, we quantified the absolute difference of the translation status per gene between syngas and fructose (TL-STATUS). Genes were then ranked by decreasing order of the calculated TL-STATUS values for evaluation with the CONSENSUS approach (see below).

Second, as the TL-PROFILE metric, we calculated for each gene the difference in its polysome profiles between syngas and fructose as the variance-weighted Aitchison’s distance of the centered-ln-ratio transformed compositional data by weighing the difference between the individual fractions by pooled variance across syngas and fructose bio-replicates. Genes were then ranked by decreasing order of the calculated TL-PROFILE values for evaluation with the CONSENSUS approach.

Third, as the POLY-FRACTIONS metric, we estimated 95% confidence intervals (CI) for each polysome fraction and gene by bootstrapping from a resampling distribution and then identified fractions with differential mRNA proportions if their computed CI did not cross the null value. This metric quantified both the number of statistically different fractions per gene and the maximal difference across fractions between conditions. Genes were then ranked by decreasing order of both features for evaluation with the CONSENSUS approach.

Lastly, to account for the different nature of translational regulation captured by each metric, we used a CONSENSUS approach that ranked genes based on the average of the ranks obtained across the three metrics. We determined DTLs as genes within the top 5% of the CONSENSUS ranking.

### RNA-seq

#### Sample preparation and sequencing

The same cell pellets as for polysome-seq were used for RNA-seq (see above for sampling details). Total RNA from frozen samples was extracted using the RNeasy Midi Kit (Qiagen) combined with on-column DNase I treatment. RNA amount was quantified using the Nanodrop spectrophotometer and RNA integrity validated using the Agilent Bioanalyzer 2100. External ERCC RNA spikes were added to 1 µg of each total RNA sample according to manufacturer instructions. Total RNA was ribo-depleted using the RIBO-Zero kit and size distribution evaluated with the Agilent Bioanalyzer 2100. After fragmentation, the first strand cDNA was synthesized using random hexamer primers. During the second strand cDNA synthesis, dUTPs were replaced with dTTPs in the reaction buffer. The directional library was ready after end repair, A-tailing, adapter ligation, size selection, USER enzyme digestion, amplification, and purification. Finally, libraries were sequenced using the Illumina Novaseq X Plus Series platform with 2 x 150 bp cycles (paired-end).

#### Quantification of transcript abundances and differential expression analysis

The ERCC RNA spikes were used to determine the linearity range between read counts and RNA concentration and thus transcripts with <10 read counts were excluded for reliable quantification. Additionally, reads with >10% of undetermined bases (N) and reads with >50% of bases with Qscore ≤ 5 were excluded. Reads trimmed of Illumina adapters were then mapped to the genome of C. autoethanogenum strain LAbrini (GenBank CP110420.1) and ERCC RNA spikes using Bowtie2^134^ and read counting was performed using the featureCounts function within Rsubread^135^. Raw library sizes were normalized across the samples using the *normLibSizes* function within edgeR^138^. Differential expression analysis between syngas and fructose cultures was performed within edgeR using the quasi-likelihood F-test as per the *glmQLFTest* function^139^. To determine genes with statistically different expression (q-value < 0.05), we applied the Benjamini–Hochberg procedure to the F-test results controlling the false discovery rate (FDR) at 5%^133^.

### Identification of RNA-binding proteins

Potential RNA-binding proteins in *C. autoethanogenum* were identified using PRONTO-TK (10.1093/nargab/lqae112), a neural network-based framework for protein function prediction. Proteins predicted with posterior probabilities above the confidence threshold of 0.75 were retained as candidate RBPs for downstream analyses of RBP–RNA interactions.

### Prediction of protein-RNA interactions

We employed catRAPID omics v2^51^ to predict protein–RNA interactions between the candidate RNA-binding proteins and the genes which turned out to be differently regulated in transcription and translation between growth conditions. The rating system of catRAPID predictions integrates multiple scores to provide final scores ranging in the interval [0,1]. Protein-RNA interactions predicted by catRAPID were retained if the corresponding Z-scores pertaining to the interaction propensity and the final scores were ranked in the top 25% positions.

## Data availability

All raw sequencing datasets have been deposited at the European Nucleotide Archive under project accession code PRJEB102336. Source data is provided in this paper.

## Acknowledgements

This work was funded by the National Recovery and Resilience Plan (NRRP), Mission 4 Component 2 Investment 1.3 - Call for tender No. 1561 of 11.10.2022 of Ministero dell’Universita’ e della Ricerca (MUR), funded by the European Union – NextGenerationEU. The project is identified by Project code PE0000021, Concession Decree No. 1561 of 11.10.2022 adopted by Ministero dell’Universita’ e della Ricerca (MUR), CUP – E13C22001890001, according to attachment E of Decree No. 1561/2022, Project title ‘‘Network 4 Energy Sustainable Transition – NEST (Spoke 3)’’. Furthermore, this work was funded by the European Union’s Horizon 2020 research and innovation program under grant agreement N810755.

## Author contributions

Conceptualization: A.R. and K.V.; Methodology: A.R., K.R., A.B., and K.V.; Formal analysis: A.R., G.M.M.P., A.B., L.G., M.C-B., and K.V.; Investigation: A.R., G.M.M.P., K.R., L.G., M.C-B., and K.V.; Software: A.R., G.M.M.P., A.B.; Resources: A.R. and K.V.; Writing - Original Draft: A.R. and K.V.; Writing - Review & Editing: A.R., K.R., A.B., L.G., M.C-B., and K.V.; Visualization: A.R and K.V.; Supervision: A.R., M.C-B. and K.V.; Project Administration: A.R and K.V.; Funding acquisition: A.R and K.V..

