## Supplementary Figures for "Systems-level analysis identifies protein-RNA interactions contributing to autotrophy of *Clostridium autoethanogenum*"

Re *et al.*

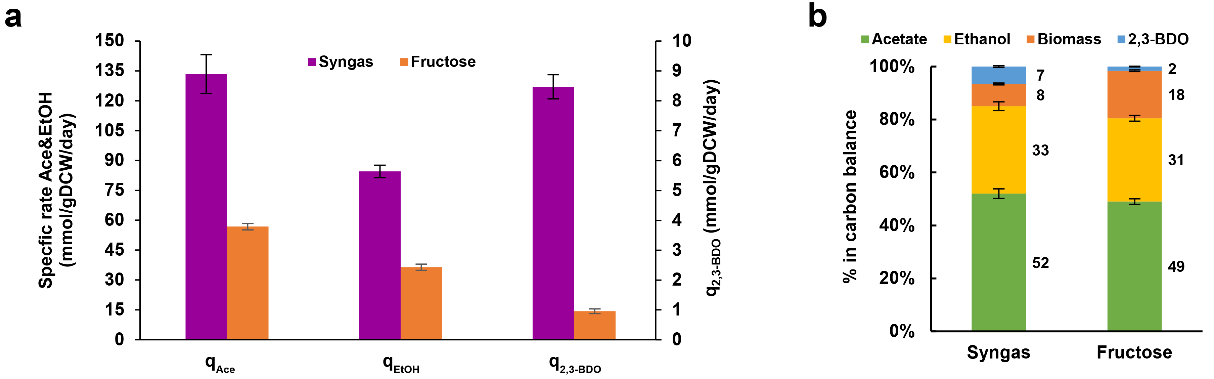

**Supplementary Figure 1**. **Specific by-product production rates and carbon balances in *C. autoethanogenum*** **syngas and fructose chemostats. a** Specific by-product production rates (mmol per gram of dry cell weight per day). Ace, acetate; EtOH, ethanol; 2,3-BDO, 2,3-butanediol. **b** Carbon balances. Carbon recoveries were normalised to 100% to have a fair comparison of carbon distributions between conditions. Bars show average ± standard deviation between biological triplicate chemostat cultures.

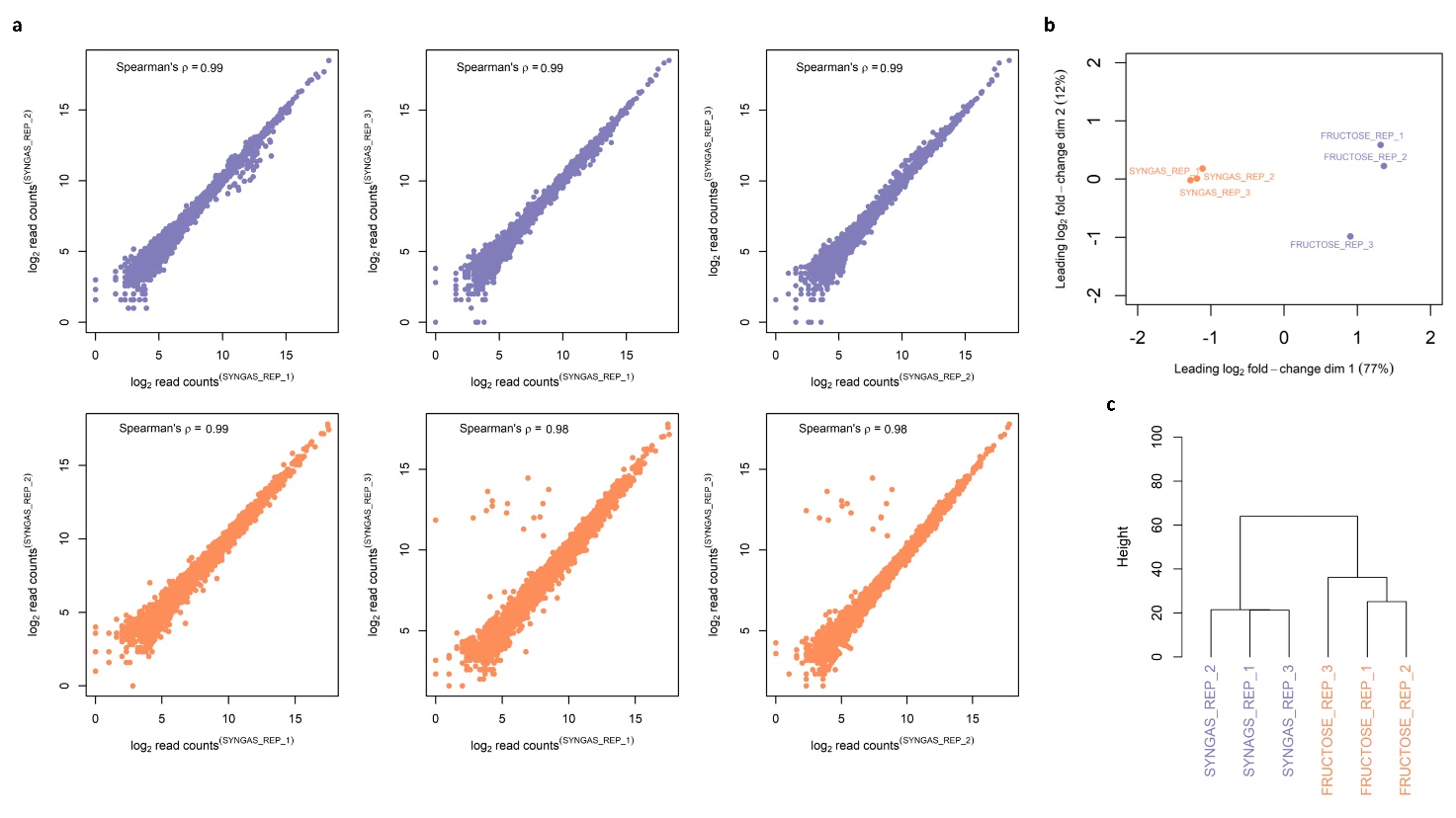

**Supplementary Figure 2**. **Reproducibility and clustering of RNA-seq data across *C. autoethanogenum* syngas and fructose chemostats. a** Correlation of normalized log_2_ RNA-seq read counts between syngas and fructose biological triplicate chemostat cultures. **b** Clustering of syngas and fructose biological triplicate chemostat cultures on a multidimensional scaling plot. Distances represent leading log_2_ fold-changes that are the root-mean-square average of the top 500 largest log_2_ fold-changes between any two samples. **c** Clustering of syngas and fructose biological triplicate chemostat cultures on a dendrogram based on normalized log2 RNA-seq read counts.

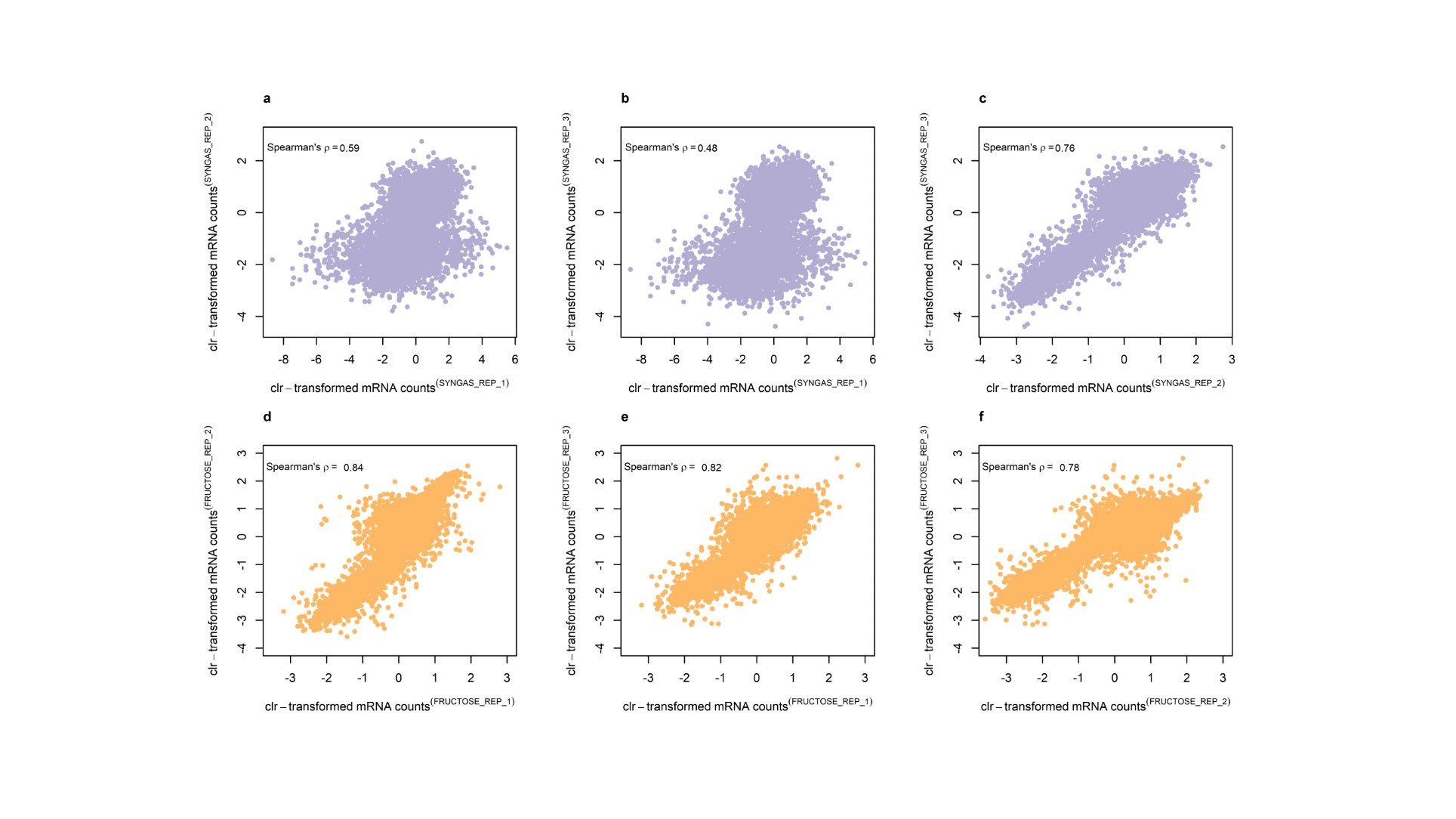

**Supplementary Figure 3.** **Reproducibility of polysome-seq data across *C. autoethanogenum* syngas and fructose chemostats.** **a-c** Correlation of centered-ln-ratio transformed (clr) mRNA counts per polysomal gradient fraction from total counts between biological triplicate syngas chemostat cultures. **d-f** Correlation of clr mRNA counts per polysomal gradient fractions from total counts between biological triplicate fructose chemostat cultures.

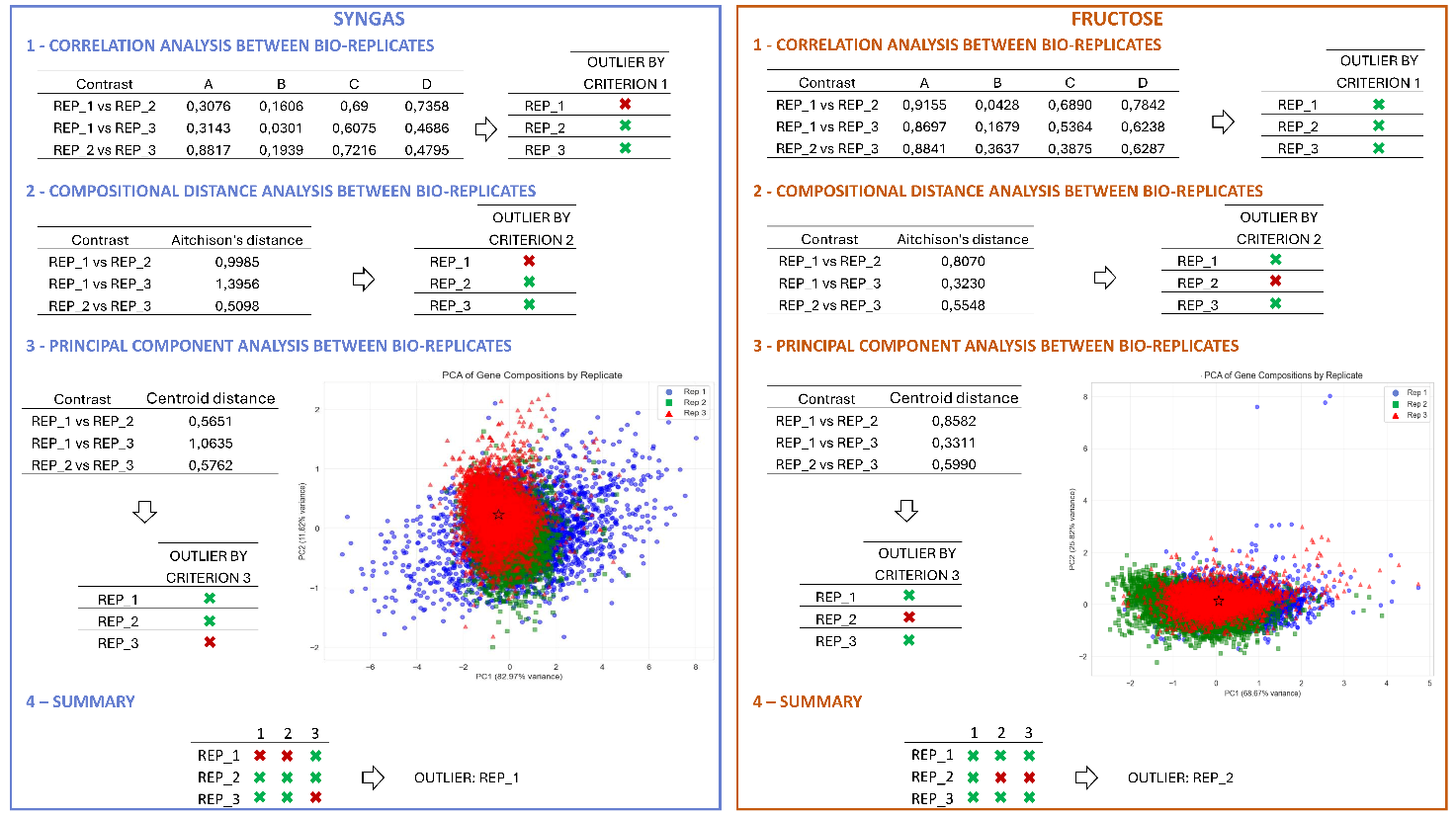

**Supplementary Figure 4. Outlier detection in polysome-seq data across bio-replicate *C. autoethanogenum* syngas and fructose chemostats.** Consistency between bio-replicates within both chemostat datasets was assessed by three complementary methods: 1) Correlation analysis of mRNA counts per polysomal gradient fractions from total counts between bio-replicates by Spearman’s rank correlation coefficient $\rho$. A bio-replicate was determined as an outlier if it showed the highest number of poorly correlated polysomal fractions; a fraction was considered poorly correlated if its data correlated poorly (ρ < 0.7) with the other bio-replicates which in turn correlated between each other with ρ > 0.7. 2) Compositional distance analysis by first calculating the average of all mRNA counts for each fraction and bio-replicate, then centered-ln-ratio transforming the averages, and lastly computing the Aitchison’s distance between each pair of bio-replicates. A bio-replicate was determined as an outlier if it showed the maximal average Aitchison’s distance from the other bio-replicates that was greater than the median distance observed across syngas and fructose. 3) Principal component analysis (PCA) by first collating all mRNA counts for each fraction and bio-replicate as a concatenated dataset, then centered-ln-ratio transforming gene compositional data and lastly performing PCA to compute the average distances between the centroids of each pair of bio-replicates. A bio-replicate was determined as an outlier if its centroid showed the maximal average distance from the centroids of the other bio-replicates that was greater than the median distance observed across syngas and fructose.

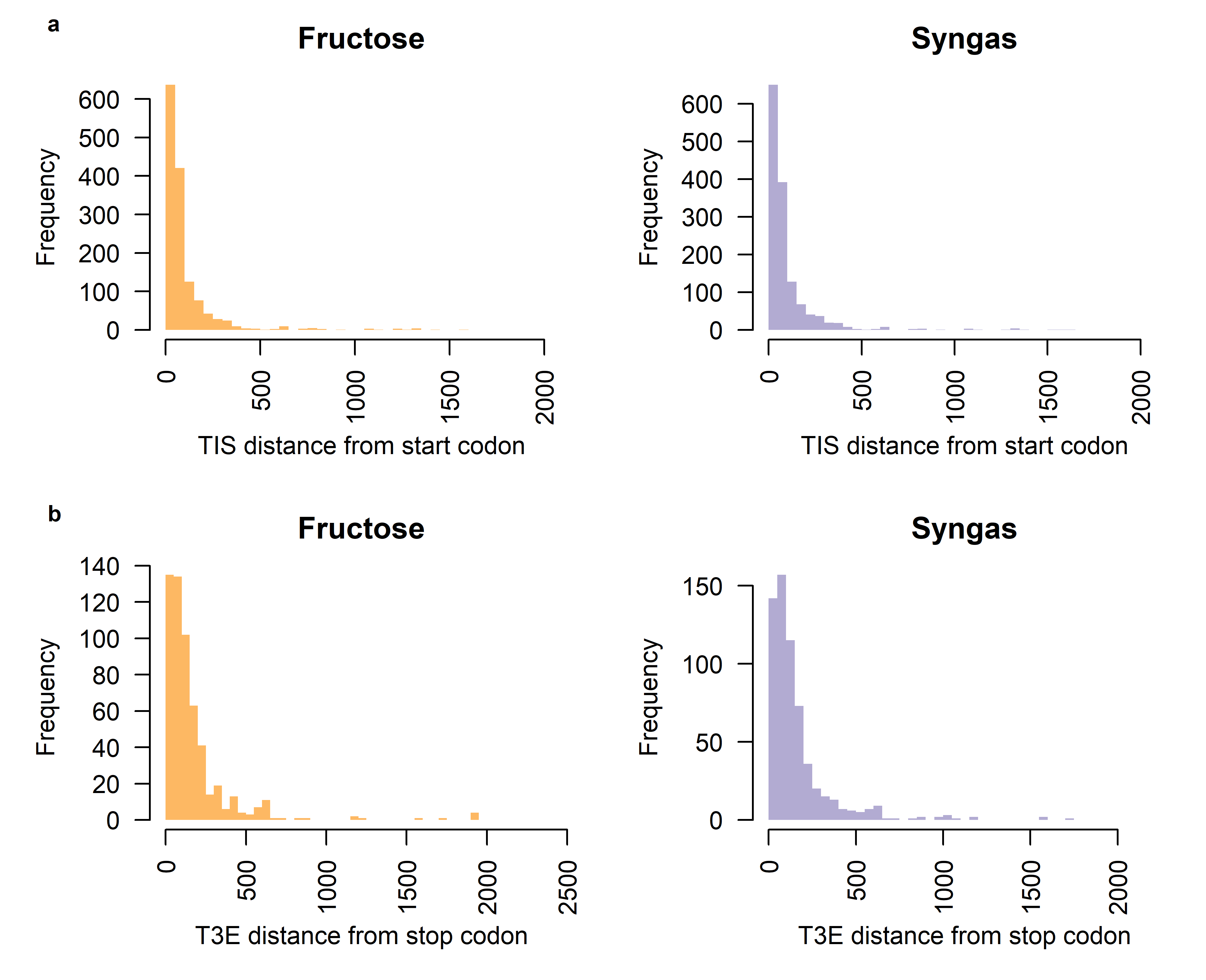

**Supplementary Figure 5**. **Distribution of 5’UTR and 3’UTR lengths in *C. autoethanogenum* syngas and fructose chemostats.** **a** 5’UTR length as distance of transcription initiation site (TIS) from start codon of coding DNA sequence. **b** 3’UTR length as distance of transcription 3’ end (T3E) from stop codon of coding DNA sequence.

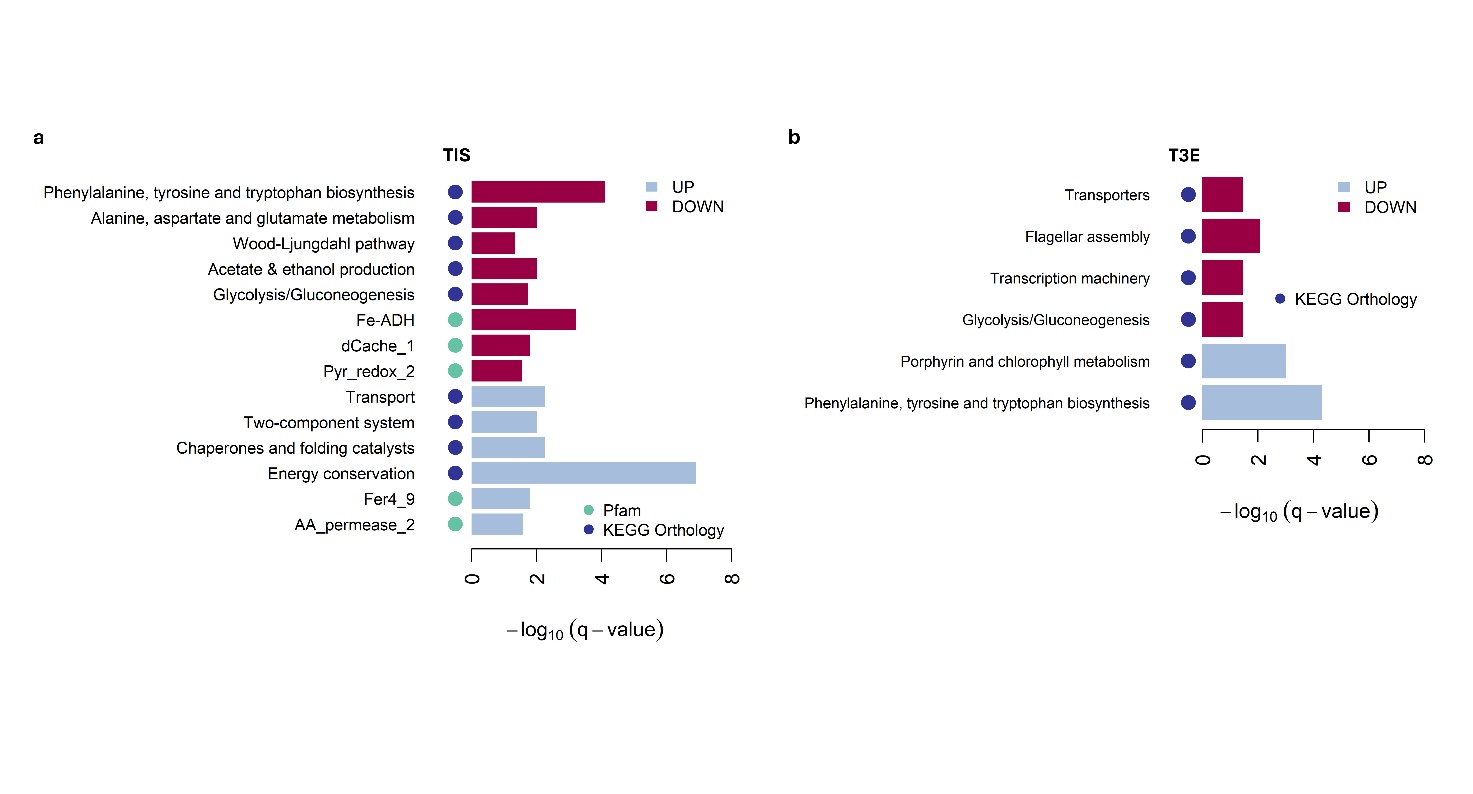

**Supplementary Figure 6**. **Functional enrichment of genes featuring differentially quantified TISs and T3Es**. **a** The barplot shows the functional classes enriched in genes featuring differentially quantified TISs by hypergeometric test (FDR = 5%). Pfam, Pfam entries (10.1093/nar/gkaa913) from eggNOG-mapper 2.1.12 (10.1093/molbev/msab293); KEGG Orthology, KEGG Orthology (KO) functional categories (10.1016/j.jmb.2015.11.006) from Valgepea et al. (Level 3 in Table S3; 10.1128/msystems.00026-22). **b** The barplot shows the functional classes enriched in genes featuring differentially quantified TISs by hypergeometric test (FDR = 5%).

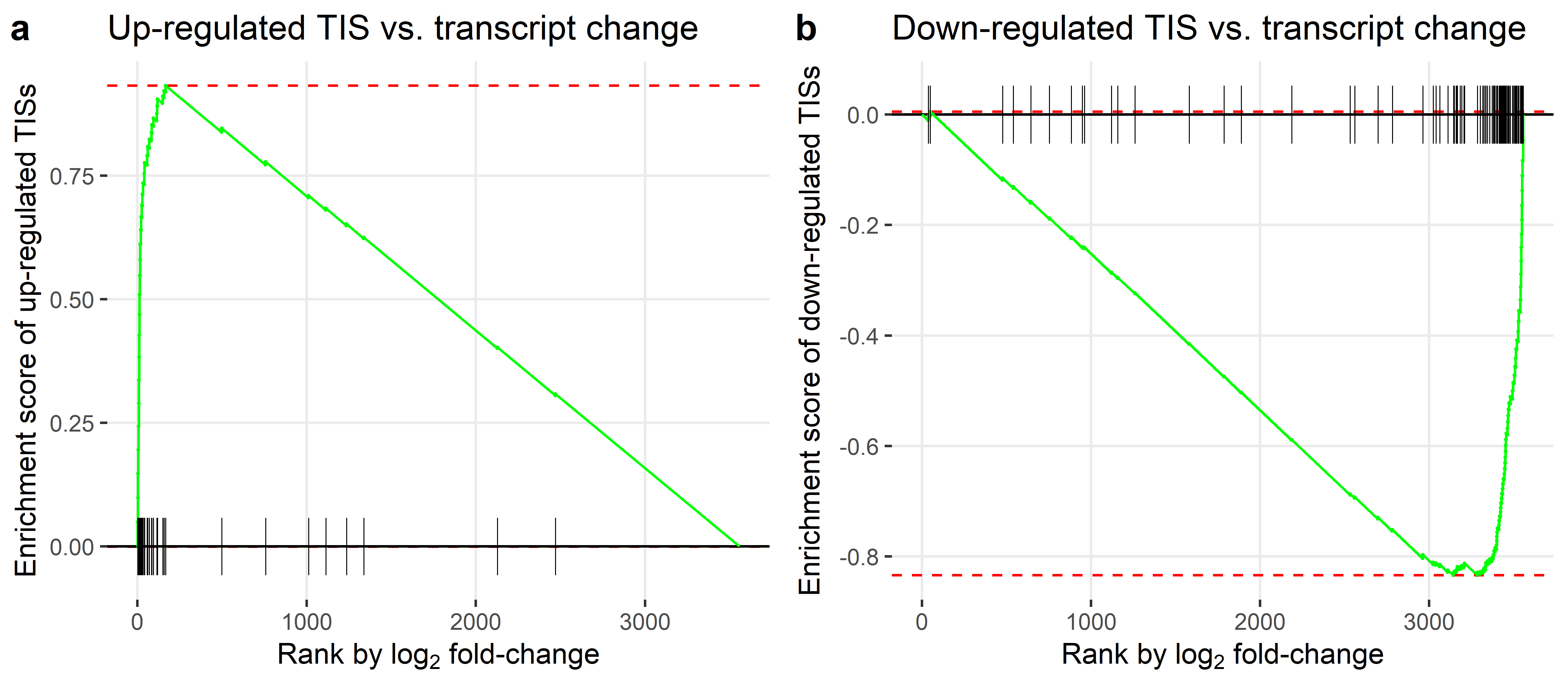

**Supplementary Figure 7**. **Enrichment of genes with differential transcription initiation signals within top or bottom ranks of transcript expression changes between syngas vs fructose *C. autoethanogenum* chemostats.** Black lines indicate the location of genes with differential signals of transcription initiation while red lines indicate the upper and lower bounds of the enrichment score. TIS, transcription initiation site.

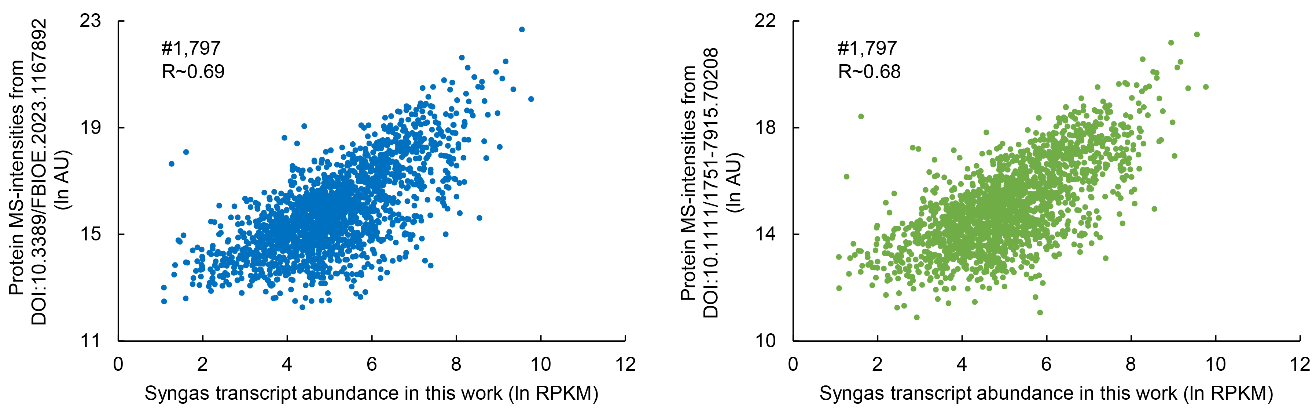

**Supplementary Figure 8**. **Correlation between transcript levels quantified in this work for *C. autoethanogenum* syngas chemostats and protein levels quantified in previous works for *C. autoethanogenum* syngas chemostats.** All datasets from *C. autoethanogenum* strain LAbrini chemostats operated at pH 5, 37°C, and dilution rate 1 day^-1^. R, Pearson product correlation; RPKM, reads per kilobase of transcript per million mapped reads; MS, mass spectrometry.
